# SAF-A/HNRNPU regulates euchromatin dynamics and is required for nuclear envelope integrity

**DOI:** 10.64898/2026.09.10.750682

**Authors:** Kaitlyn Alimenti, Matteo Mazzocca, Anders S Hansen, Michael D Blower

## Abstract

Defects in nuclear morphology are associated with cancer and premature ageing. Nuclear morphology is established by a balance of external forces produced by actin contractility and resistance provided by chromatin and the nuclear lamina. Euchromatin is decompacted to facilitate transcription, which has been shown to decrease local nucleosome motion. However, the role of transcription and euchromatic chromatin structure in nuclear envelope integrity are not clear. Here, we demonstrate that acute depletion of SAF-A/hnRNPU causes compaction of euchromatic regions and constriction of chromatin mobility, leading to loss of nuclear envelope integrity in a transcription-dependent manner. SAF-A is required in a dose-dependent manner to preserve nuclear shape under conditions of nuclear softening and elevated transcription. Our work identifies SAF-A as a dynamic scaffold that regulates chromatin structure and dynamics at sites of active transcription to preserve nuclear envelope mechanical tension.

## Introduction

Maintenance of nuclear architecture is essential for proper gene expression and cellular fitness. Aberrant nuclear shape is a hallmark of genomic instability, cancer, and age-related disorders^1,2^. Nuclear morphology is maintained by a force balance between two major components: 1) cytoskeletal tension exerted on the nucleus and 2) mechanical resistance from the nuclear envelope, lamins, and chromatin. Decreasing the global heterochromatin-to-euchromatin ratio by histone deacetylase inhibition or depletion of histone modifying enzymes leads to changes in nuclear stiffness and defects in nuclear shape^3–7^. It is unclear if loss of heterochromatin is the sole driver of these defects or if euchromatin may also play an active role in abnormal nuclear morphology development.

Euchromatin is decondensed and enriched for actively transcribed genes. Active RNA polymerase II (RNAPII) locally concentrates into “hubs” containing nascent transcripts and associated proteins^8,9^. Regions of active transcription require dramatic chromatin remodeling as the chromatin fiber is decompacted, stretched, and subjected to mechanical forces generated by RNA polymerase II (RNAPII) motor activity^10–12^. At transcription hubs, tracking of single nucleosomes revealed that transcription decreases local chromatin motion^13–15^. Recent studies suggest that transcription contributes to nuclear envelope mechanical tension, as decreasing transcriptional activity rescues nuclear shape defects in cells with lamin or chromatin perturbations^16,17^. How euchromatic regions coordinate transcription-associated forces, and whether transcriptional hubs provide mechanical rigidity to the nucleus is not known.

Scaffold attachment factor A (SAF-A), also known as heterogeneous nuclear ribonucleoprotein U (hnRNPU), is a highly abundant nuclear RNA binding protein required for transcriptional regulation, mRNA splicing, and long noncoding RNA (lncRNA) localization to chromatin. Loss of SAF-A is lethal in human tissue culture cells and embryonically lethal in mice^18–20^. Haploinsufficiency in humans causes a neurodevelopmental disorder (HNRNPU-NDD) with hallmark phenotypes of developmental delays, seizures, and cardiac abnormalities^21–23^.

SAF-A was previously proposed to be a regulator of euchromatin organization. SAF-A knockdown leads to more condensed euchromatic regions in primed stem cells, fully differentiated human cell lines, and mouse cells^24–26^. A recent preprint suggests that binding of SAF-A to chromatin associated RNAs (caRNAs) forms microphase separated meshworks at sites of RNAPII activity that regulate nuclear fluid dynamics^27^. SAF-A may influence chromatin organization through transient interactions with nascent transcripts and DNA to create a dynamic nuclear scaffold at sites of active transcription. Consistent with this model, SAF-A is a highly dynamic protein, with a rapid Fluorescence Recovery After Photobleaching (FRAP) recovery half-time of ∼2.5 seconds^27,28^. Transcriptional inhibition dramatically increases SAF-A nuclear mobility, consistent with SAF-A dynamically binding to nascent RNA^28,29^. Therefore, SAF-A:RNA complexes may be critical to providing structural integrity to sites of active transcription.

Here, we employed an auxin-inducible degron (AID) system to study the role of SAF-A in nuclear morphology and nuclear envelope integrity. We find that acute depletion of SAF-A causes severe nuclear morphology defects, nuclear envelope rupture, and loss of chromatin mobility. We show that SAF-A is a concentration-dependent structural scaffold critical for euchromatic architecture and nuclear envelope integrity. Together, these findings suggest that SAF-A forms a dynamic molecular meshwork at sites of active transcription to regulate local chromatin dynamics. Our work establishes a mechanistic link between transcriptional activity and nuclear morphology, revealing a novel role of euchromatin in regulating global nuclear architecture.

## Results

### SAF-A depletion leads to abnormal nuclear morphology

To investigate whether SAF-A is required to maintain nuclear structure, we used an auxin-inducible degron system previously established in diploid, karyotypically-stable RPE-1 cells^28–30^ (Figure S1A). Simultaneous treatment with 1 µM doxycycline (dox) and 1 µM indole-3-acetic acid (IAA) for 24 hours lead to near-complete depletion of SAF-A-mAID-mCherry (Figure S1B)^28,29^. SAF-A-mAID-mCherry RPE-1 cells were treated with IAA and doxycycline for 24, 48, and 72 hours, fixed, and imaged by confocal microscopy to evaluate interphase nuclear shape. Abnormal nuclear morphology was defined by the presence of polylobulations, nuclear blebs, or micronuclei^31–33^. A subset of cells (∼10%) exhibited abnormal nuclear morphology as early as 24 hours post-SAF-A depletion and increased progressively over time, with >20% displaying nuclear defects by 72 hours (Figure S1C). To determine if nuclear morphology defects are also present in live cells, we integrated YFP-H2B into our SAF-A-mAID-mCherry RPE-1 cells via lentiviral transduction. Consistent with fixed cell analysis, a subset of 24-hour SAF-A depleted cells (∼10%) exhibited abnormal nuclear morphology, which progressively increased over the three-day time course (Fig. 1AB). These results confirm that acute SAF-A depletion is sufficient to cause nuclear morphology defects within 24 hours, and that the affected population expands over time.

**Figure 1.**
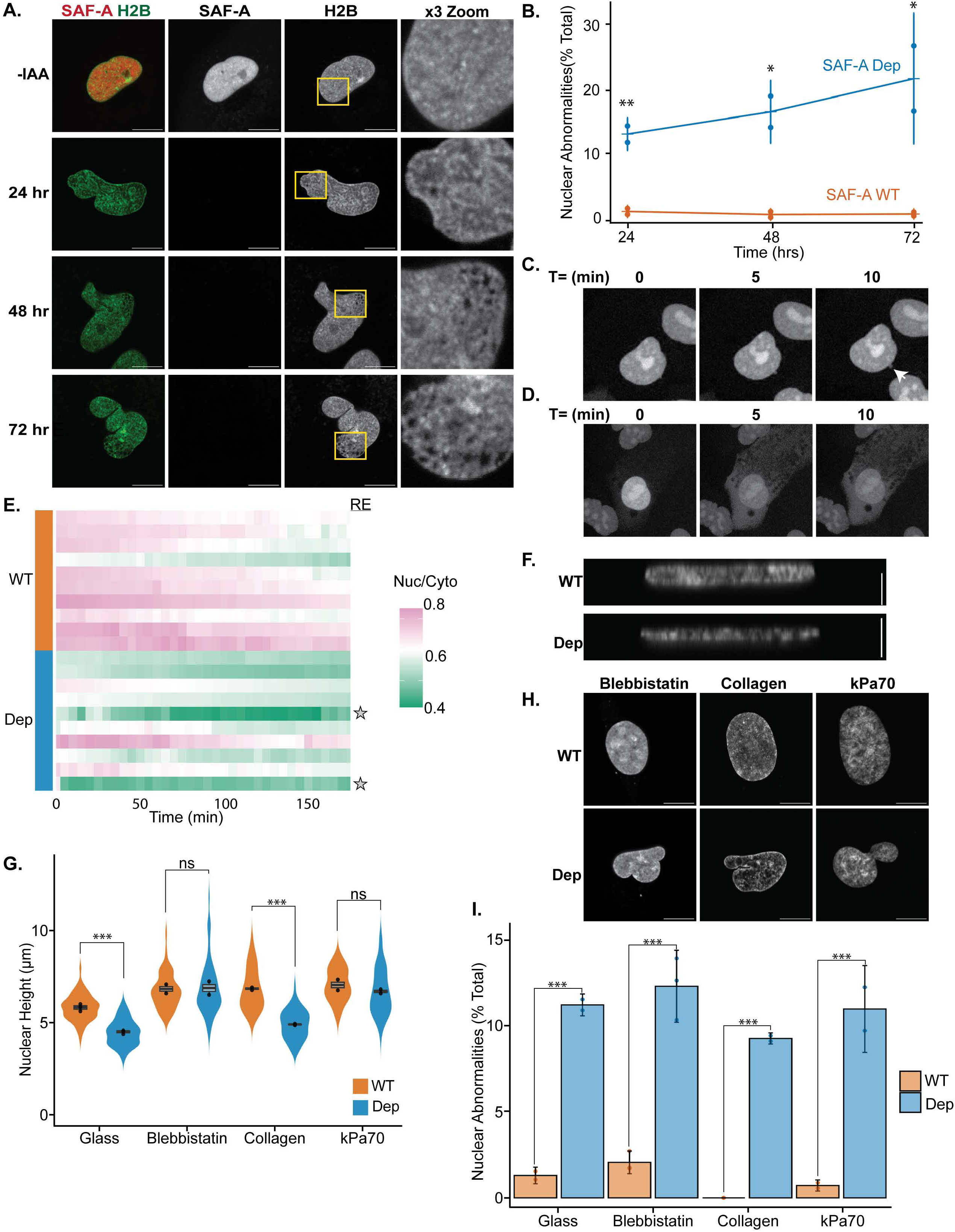
**A.** Fluorescent imaging of SAF-A-mAID-mCherry and YFP-H2B in live RPE1 cells treated with IAA and doxycycline for 24, 48, or 72 hrs. **B.** Scoring of nuclear morphology defects from cells imaged in A. Statistical comparison of percent nuclear abnormalities was conducted using a one-way ANOVA followed by Tukey’s test with a Bonferroni correction. The p value comparisons of -IAA and +IAA were: 24 hr p= 0.02, 48 hr p= 0.023, 72 hr p= 0.053. **C.** Representative images of formation of nuclear bleb in cells expressing NLS-HALO. Images are rendered as maximum projections of a 3D stack taken in 5-minute intervals. **D.** Representative images of formation nuclear envelope rupture event. **E.** Quantification of the ratio of nuclear to cytoplasmic NLS-HALO fluorescence intensity over a 3-hour live cell imaging time course. Ten cells were monitored for SAF-A wildtype and SAF-A depletion conditions. Cells with a nuclear rupture (RE) events are denoted with a gray star. **F.** DAPI staining of SAF-A-mCherry AID RPE-1 cells. Images are rendered as an orthogonal view of a 3D stack. Bar, 5 µM. **G.** Quantification of nuclear height in WT and SAF-A depleted cells with blebbistatin co-treatment or seeded on glass or collagen or Kpa70 polyacrylamide hydrogels surface. Statistical comparison was conducted by one-way ANOVA with a Bonferroni correction. The p value comparisons of SAF-A WT glass vs. collagen p=0.002, SAF-A WT glass vs. kPa70 p=0.0004, SAF-A Dep glass vs. collagen p=0.3, SAF-A Dep glass vs. kPa70 p=0.000002. **H.** DAPI staining of SAF-A-mAID-mCherry RPE-1 cells seeded on glass, collagen or kPa70 polyacrylamide hydrogels. The glass seeding condition was treated with blebbistatin. Images are rendered as a maximum projection of a 3D stack. Bar, 10 µM. **I.** Scoring of nuclear abnormalities in experimental conditions from G. and H. Data from two biological replicates was graphed as percent of cell population with nuclear abnormalities. Statistical comparison was conducted by one-way ANOVA with a Bonferroni correction. The p value comparisons are glass SAF-A WT vs. SAF-A Dep p=0.0001, collagen1 SAF-A WT vs. SAF-A Dep p=0.0002, kPa70 SAF-A WT vs. SAF-A Dep p=0.0001. Cell and replicate n are listed in Materials and Methods.

A subset of SAF-A depleted cells in fixed and live image analysis exhibited pronounced nuclear blebs, well-characterized sites of transient nuclear envelope rupture. To analyze nuclear envelope integrity we generated a reporter cell line by stably integrating a doxycycline-inducible NLS-HaloTag construct into the SAF-A-mAID-mCherry background via lentiviral transduction^34,35^(Figure S1DE). Cells labeled with Oregon Green HaloTag dye following SAF-A depletion were monitored using live cell imaging to assess nuclear compartmentalization. Over a 3 hour imaging period ∼25% of the cells monitored exhibited abnormal nuclear morphology (Fig. 1C). We did not observe repair of nuclear envelope defects during the course of imaging. In addition, 20% of the cells imaged experienced a nuclear rupture, none of which were able to re-establish nuclear compartmentalization (Fig. 1DE, Figure S1F). Nuclear ruptures were defined as local minima in the nuclear-to-cytoplasmic GFP ratio corresponding to a ≥20% reduction from the initial ratio. Our live cell imaging analysis revealed that SAF-A is required to maintain nuclear compartmentalization, and its acute depletion results in compromised nuclear envelope integrity.

Previous work has shown that SAF-A depletion leads to immediate changes in mRNA splicing and global changes to gene expression following depletion for 48 or 72 hours^25,28,29^. To determine if SAF-A depletion leads to altered splicing or expression of proteins important for nuclear morphology we examined RNA-seq data of nuclear pore components and lamins from cells depleted of SAF-A for 24 hours. Splicing and expression of nuclear pore components and lamins were not affected in SAF-A Dep cells (Figure S2AB). Western blot analysis of 24 hour SAF-A depleted cells showed no quantitative difference in Lamin A/C or Lamin B1 expression (Figure S2C-F). These results suggest that SAF-A plays a direct role in the regulation of nuclear envelope integrity.

We recently demonstrated that SAF-A depletion leads to cell cycle exit over a 10-day time course^28^. To determine if SAF-A depleted cells with nuclear abnormalities were actively dividing we performed immunofluorescence for Mitosin/CENP-F, a kinetochore associated protein with tightly cell-cycle regulated expression^36^. Notably, SAF-A depleted cells with aberrant nuclear morphology exhibited a significant reduction of nuclear Mitosin intensity compared to morphologically normal cells across the 3-day time course (Figure S3AB). In addition, cells with abnormal nuclear morphology following 24 hours of SAF-A depletion showed a significant increase in p21 expression compared to cells with normal morphology, suggesting abnormal SAF-A depleted cells are arrested in G1 (Figure S3CD). A recent publication demonstrated that altered nuclear membrane tension caused by chromosome missegregation elevated p21 levels^2^ and that two mTORC2 inhibitors (INK-128 and JR-AB2-011) decreased p21 levels in cells under nuclear envelope stress. To test whether mTORC2 inhibition could restore nuclear morphology in SAF-A–depletion cells, we treated SAF-A Dep cells with INK-128 or JR-AB2-011 for 24 hours. SAF-A depleted cells treated with mTORC2 inhibitors showed no reduction in nuclear morphology defects compared to the DMSO control group (Figure S3C,E). Collectively, these findings demonstrate that SAF-A depletion leads to progressive nuclear morphology defects in cells that accumulate in G1 and triggers a p21-mediated nuclear envelope stress pathway.

### SAF-A depleted cells have increased actin-based nuclear confinement

We hypothesized that morphology defects following SAF-A depletion may be due to decreased resistance to force exerted on the nucleus by actin. To evaluate the ability of SAF-A depleted cells to resist the mechanical tension of actin contractility we measured nuclear height of RPE-1 cells depleted of SAF-A for 24 hours. SAF-A depleted cells exhibited a ∼1.5 µm decrease in nuclear height (Fig. 1FG) consistent with nuclear softening and a decreased capacity to provide rigidity against actin contractility

To evaluate if actin confinement is a driver of nuclear morphology defects in SAF-A depleted cells, we perturbed actin contractility by two complementary approaches. First, we altered actin organization by softening cell seeding surfaces, as actin polymerization is dependent on substate rigidity (Doss et al., 2020). SAF-A-mAID-mCherry cells were seeded on collagen 1 matrix or polyacrylamide gels with 70 kPa measured stiffness^37–39^. Following a 24-hour incubation period, cells were treated with IAA for 24 hours to deplete SAF-A. Consistent with previous results, nuclear height increased as a function of substrate stiffness^40,41^ (Figure S1G). Nuclear height was significantly reduced in SAF-A depleted cells compared to untreated controls in collagen of 70 kPa substrate conditions (Fig. 1G). SAF-A depleted cells incubated on all seeding substrates had nuclear morphology defects in ∼10% of the population with no statistically significant differences when compared to glass (Fig. 1H,I). As an orthogonal approach, we co-treated SAF-A-mAID-mCherry cells with IAA and the myosin II inhibitor blebbistatin to prevent assembly of actin bundles^42^. As expected, blebbistatin treatment increased nuclear height in DMSO control cells suggesting a reduction in cytoskeleton tension (Figure S1G). In addition, blebbistatin treatment fully rescued nuclear height defects in SAF-A depleted cells (Fig. 1G).

However, treatment of SAF-A depleted cells with blebbistatin did not rescue nuclear shape defects (Fig. 1H,I). Therefore, nuclear morphology defects following SAF-A depletion are not driven by actin-mediated force.

We considered other cellular mechanisms that may promote nuclear morphology defects in SAF-A depleted cells. Nuclear softening suggests SAF-A depleted cells may have a decrease in the global heterochromatin: euchromatin ratio. However, global fluorescence intensity levels of DAPI staining and immunofluorescence for H3k9me3, RNAPII Ser2P, and RNAPII Ser5P following 24 hour SAF-A depletion were not altered, suggesting loss of SAF-A does not cause global expansion of euchromatin or loss of heterochromatin (Figure S1H). Therefore, nuclear softening following SAF-A depletion may be caused by changes in chromatin dynamics.

### SAF-A depletion restricts nucleosome motion

SAF-A is proposed to interact with nascent transcripts to form a microgel at sites of active transcription that may affect nuclear fluidity^27^ and space euchromatic genome regions^25^ which could affect chromatin mobility. To determine if SAF-A regulates chromatin motion, we integrated a H2B-HaloTag construct into our SAF-A-mAID-mCherry RPE-1 cells and preformed single particle tracking (SPT) of tagged nucleosomes^43^. To record chromatin motion, we tracked H2B-Halo across three orders of magnitude in time, spanning the timescale over which we could previously observe changes in chromatin dynamics^43^. To overcome photobleaching and achieve this dynamic range, we recorded H2B-Halo using two time lags (100 ms and 2 seconds). To quantify chromatin motion across the full timescale, we next computed the mean squared displacement (MSD) for each time lag and condition (Figure S4). We first observed that chromatin dynamics in untreated RPE-1 are highly subdiffusive, with an MSD well described by power law behavior (α ≈ 0.3), confirming our previous finding (Figure S4A)^43^. Under SAF-A knockdown, chromatin dynamics follow a similar pattern, with a highly sub-diffusive exponent. To better quantify differences in chromatin dynamics, we computed the relative chromatin mobility which quantifies how much chromatin moves as a fold change relative to the control, allowing comparison across conditions (Fig. 2B). SAF-A depleted cells exhibited a decrease in the relative mobility of tagged nucleosomes (Fig. 2B) comparable to that observed following topoisomerase II inhibition^43^. Our results suggest that SAF-A promotes chromatin motion at sites of active transcription with its depletion potentially leading to a compaction of gene rich regions^25^, which in turn constrains chromatin motion. Interestingly, active transcription has previously been shown to restrict chromatin motion^13–15^. Therefore, SAF-A may counteract this restriction through dynamic interactions with chromatin and nascent transcripts. To test this theory, we inhibited RNAPII transcription in SAF-A-mAID-mCherry RPE-1 cells with triptolide for 24 hours in the presence or absence of auxin. Inspection of the MSD curves shows a remarkable effect of transcription inhibition, with the MSD increasing more rapidly at short timescales compared to untreated conditions (i.e. chromatin is more mobile) but reaching a plateau at long timescales (Figure S4C). This plateau strongly deviates from the power-law fit, pointing to chromatin dynamics that vary with timescale and/or confinement at a length scale of a few hundred nanometers. To quantify the overall effect of transcription inhibition on chromatin dynamics, we quantified chromatin mobility following triptolide relative to the untreated condition (Fig. 2B). As previously reported, transcription inhibition increased chromatin mobility in RPE-1^15^. Finally, co-treatment with triptolide and auxin resulted in a substantial reduction of relative chromatin mobility to near baseline levels, further suggesting SAF-A promotes chromatin mobility (Fig. 2B). This result is consistent with a model where SAF-A depletion may cause aggregation of chromatin that leads to a decrease in mobility in a transcription dependent manner.

**Figure 2.**
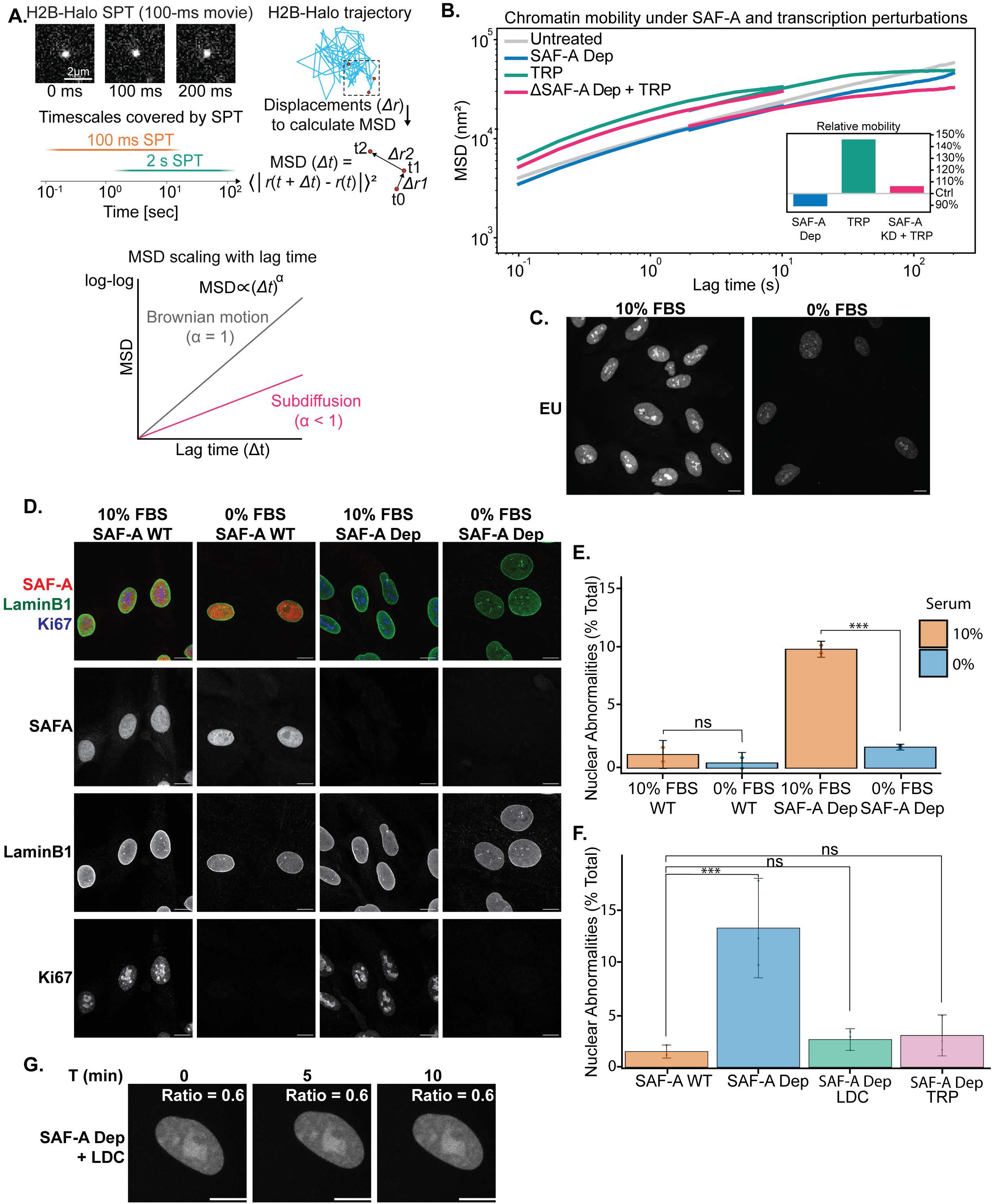
**A.** Representative frames from a 100-ms H2B-Halo SPT movie are shown together with the timescales covered by SPT, using two complementary lag times (100 ms and 2 s). H2B-Halo trajectories were reconstructed from the detected positions and used to calculate displacements (Δ*r*) over different lag times (Δ*t*). The squared displacements were then averaged at each lag time to calculate the mean squared displacement (MSD). Plotting MSD as a function of lag time gives the scaling exponent α (the MSD slope on a log-log plot), with α = 1 corresponding to Brownian motion and α < 1 to subdiffusion. **B.** Ensemble mean squared displacements (MSDs) of H2B-Halo in RPE-1 cells across treatment conditions over three biological replicates. H2B-Halo trajectories were collected using two time lags (100 ms and 2 s), and MSDs from both were integrated into a single dynamic range. Untreated (gray; 100 ms n_cells_ = 59, n_tracks_ = 3138; 2 s n_cells_ = 57, n_tracks_ = 3103), SAF-A knockdown (SAF-A KD, blue; 100 ms n_cells_ = 46, n_tracks_ = 2461; 2 s n_cells_ = 48, n_tracks_ = 2888), triptolide (TRP, teal; 100 ms n_cells_ = 58, n_tracks_ = 5815; 2 s n_cells_ = 54, n_tracks_ = 4267), and combined SAF-A knockdown and triptolide (SAF-A KD + TRP, magenta; 100 ms n_cells_ = 50, n_tracks_ = 4091; 2 s n_cells_ = 49, n_tracks_ = 4042). Average mobility relative to untreated control is shown in the inset at the bottom right. **C.** Cy3-5-EU labeled nascent transcripts in RPE1 cells cultured in 10% or 0% FBS. Images are rendered as maximum projections of a 3D stack. Bar, 10 µM. **D.** Immunofluorescence for LaminB1 and Ki67 in SAF-A depleted cells cultured in 10% or 0% FBS. Images are rendered as a maximum projection of a 3D stack. Bar, 10 µM. **E.** Quantitation of nuclear abnormalities in experimental conditions from D. Data from two biological replicates was graphed as percent of cell population with nuclear abnormalities. Statistical comparison was conducted by one-way ANOVA with a Bonferroni correction. The p value comparisons are 10% FBS SAF-A WT vs. 0% FBS SAF-A WT p=0.9 and 10% FBS SAF-A Dep vs 0% FBS SAF-A Dep p=0.0005. **F**. Quantitation of nuclear abnormalities of SAF-A depleted cells co-treated with LDC or TRP overnight. Data from three biological replicates was graphed as percent of cell population with nuclear abnormalities. Statistical comparison was conducted by one-way ANOVA with a Bonferroni correction. The p value comparisons are SAF-A WT vs. SAF-A Dep p=0.001, SAF-A WT vs. SAF-A Dep LDC p=0.002, and SAF-A WT vs. SAF-A Dep TRP p=0.003 **G.** Fluorescence imaging of labeled NLS-HALO construct in SAF-A depleted cells co-treated with LDC overnight. Live imaging was taken over three hours with 5-minute intervals. Images are rendered as maximum projections of a 3D image. Bar, 10 µM. Ratio of nuclear to cytoplasmic NLS-HALO signal presented in top right corner.

### SAF-A nuclear morphology defects are transcription dependent

To determine if active transcription is required for nuclear morphology defects following SAF-A depletion we decreased global transcriptional activity by serum starvation. RPE-1 cells cultured in 0% FBS media for 48 hours were devoid of Ki-67 expression and exhibited a 50% reduction in global transcription, indicating exit from the cell cycle (Figure 2CD, Figure S5A-C). Serum-starved SAF-A depleted cells displayed minimal nuclear shape defects, with no statistically significant differences in nuclear morphology defects compared to untreated control cells cultured in either 10% or 0% FBS (Fig. 2D-E). To directly test if transcription is a driver of nuclear morphology defects following SAF-A depletion, we co-treated SAF-A-mAID-mCherry cells with IAA and a CDK9 inhibitor that blocks transcription elongation (LDC000067, LDC) or an XBP inhibitor that blocks initiation (Triptolide, TRP) overnight^44,45^. Transcription inhibition by LDC or TRP overnight treatment resulted in a >50% decrease in nascent transcription levels compared to untreated control cells measured by EU-labeling (Figure S5D-E). SAF-A depleted cells co-treated with either inhibitor had nuclear morphology equivalent to that of untreated control cells (Fig. 2F). Therefore, we hypothesized transcription inhibition would also rescue nuclear envelope ruptures observed in our SAF-A depleted NLS-HALO reporter cell line. Cells expressing NLS-HALO for 24 hours were co-treated with IAA and LDC overnight and then monitored for loss of nuclear compartmentalization by live cell imaging. None of the SAF-A depleted cells treated with LDC exhibited nuclear envelope ruptures or sudden loss of nuclear NLS-HALO localization (Fig. 2G, S5F). Together, these findings support the conclusion that nuclear shape abnormalities and loss of nuclear compartmentalization following SAF-A depletion are transcription dependent.

### RNA binding by SAF-A is important for normal nuclear morphology

To investigate the domains of SAF-A required for maintaining nuclear morphology, we used a panel of cell lines expressing C-terminally GFP tagged SAF-A mutant rescue constructs at near-endogenous levels in the SAF-A-mAID-mCherry background^28–30^. Rescue constructs included full length SAF-A (SAF-A-^WT^-GFP), SAF-A with the SAP DNA binding domain deleted (SAF-A-^ΔSAP^-GFP), SAF-A with the RNA binding RGG repeats deleted (SAF-A-^ΔRGG1-7^-GFP), SAF-A ATP binding deficient mutant (SAF-A^K510A^–GFP), and the SAF-A ATP hydrolysis deficient mutant (SAF-A-^D580A^-GFP) (Fig. 3 A). The panel of SAF-A-GFP tagged cell lines were treated with doxycycline and IAA for 24, 48, or 72 hours to induce endogenous SAF-A depletion and expression of the respective rescue constructs. Cells were then fixed and scored for abnormal nuclear morphology. Expression of all SAF-A mutant constructs except SAF-A-^ΔRGG1-7^-GFP were able to rescue the nuclear morphology defects for all timepoints (Fig. 3 B). This result suggests SAF-A binding to nascent RNA transcripts is required to maintain nuclear shape, and is independent of its DNA binding or ATPase function^27–29^. Therefore, we propose that SAF-A binds to nascent transcripts and forms a dynamic scaffolding meshwork required to maintain euchromatin spacing and mobility, which is in turn required for normal nuclear morphology.

**Figure 3.**
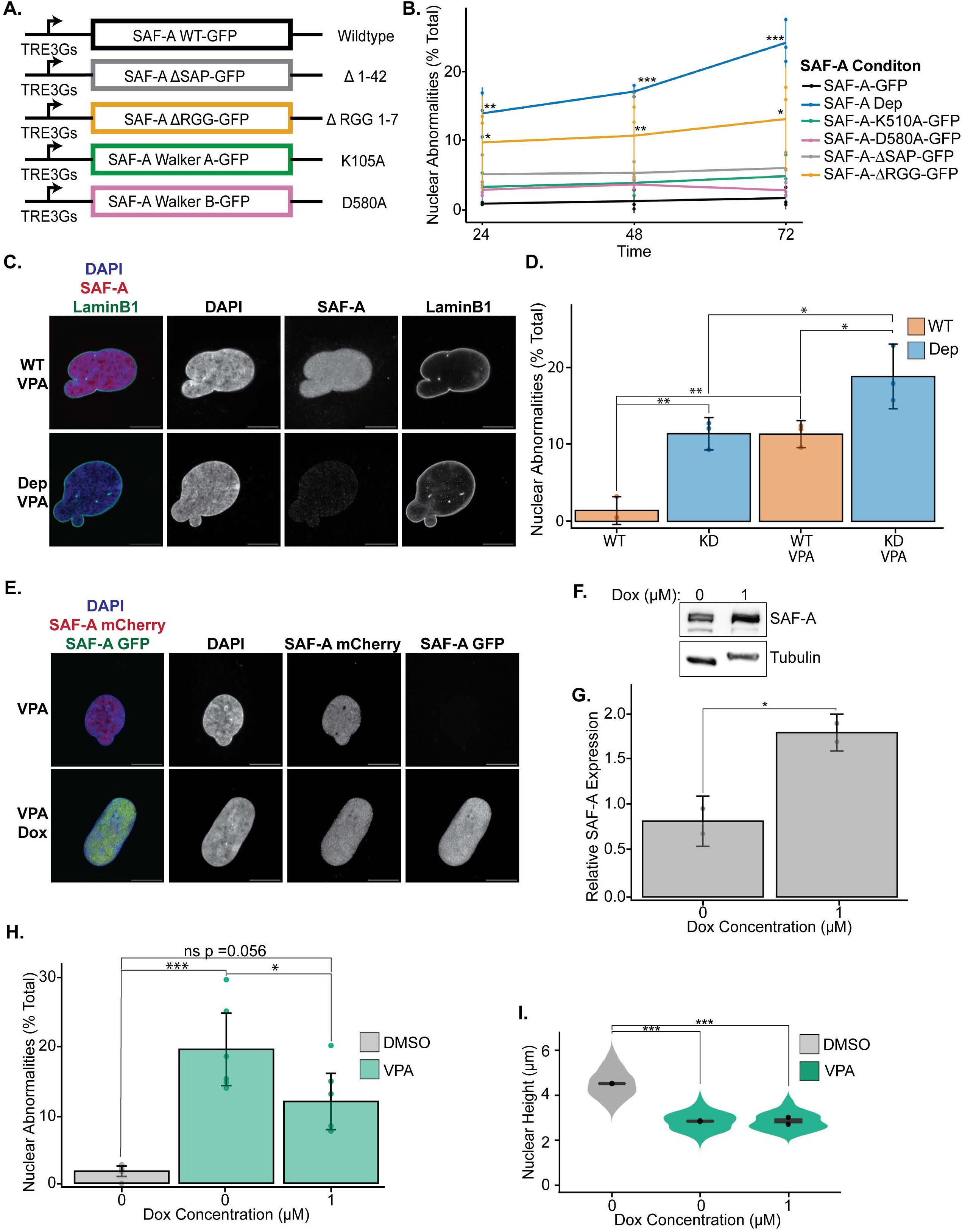
**A.** Schematic of dox inducible SAF-A rescue constructs in cells with dox inducible TiR1 and endogenous SAF-A-mAID-mCherry. **B.** Nuclear abnormalities presented as percent of total population at 24, 48, and 72 hours of SAF-A rescue construct expression. Data from three biological replicates was graphed as percent of cell population with nuclear abnormalities. Statistical comparison of percent nuclear abnormalities compared to SAF-A GFP control cells was conducted using a one-way ANOVA followed by Tukey’s test with a Bonferroni correction. The p value comparisons were: SAF-A Dep 24 hr p= 0.002, 48 hr p= 0.0001, 72 hr p= 0.00001, SAF-A K510A-GFP 24 hr p= 0.9, 48 hr p= 0.8, 72 hr p= 0.9, SAF-A D580A -GFP 24 hr p= 0.9, 48 hr p= 0.8, 72 hr p= 1.0, SAF-A ΔSAP-GFP 24 hr p= 0.5, 48 hr p= 0.4, 72 hr p= 0.6, SAF-A ΔRGG-GFP 24 hr p= 0.03, 48 hr p= 0.006, 72 hr p= 0.02. **C.** Immunofluorescence for LaminB1 in SAF-A-mAID-mCherry RPE-1 cells treated with VPA or cotreated with VPA and IAA (SAF-A Dep). Images are rendered as a maximum projection of a 3D stack. Bar, 10 µM. **D.** Quantitation of nuclear abnormalities in SAF-A-mAID-mCherry RPE-1 cells treated with VPA or cotreated with VPA and IAA (SAF-A Dep) for 24 hours. Data from three biological replicates was graphed as percent of cell population with nuclear abnormalities. Statistical comparison was conducted by one-way ANOVA with a Bonferroni correction. The p value comparisons are SAF-A WT vs SAF-A Dep p=0.003, SAF-A Dep vs SAF-A Dep VPA p=0.02, and SAF-A WT VPA vs. SAF-A Dep VPA p=0.02. **E.** Fluorescence imaging of SAF-A-mAID-mCherry RPE-1 cells with a dox inducible SAF-A-GFP rescue construct treated with VPA only or VPA and dox for 24 hours. Images are rendered as a maximum projection of a 3D stack. Bar, 10 µM. **F.** Western blot SAF-A in experimental condition outlined in E. Tubulin shown as a loading control **G.** Relative SAF-A protein expression measured by western blot analysis shown in F. The relative abundance to tubulin loading control of three biological replicates is shown as a bar graph, statistical comparison was conducted by an unpaired Student’s t test with a resulting p value of 0.04. **H.** Quantitation of nuclear abnormalities in 24-hour VPA treated SAF-A-mAID-mCherry RPE-1 cells with dox induced expression of SAF-A-GFP. Data from six biological replicates was graphed as percent of cell population with nuclear abnormalities. Statistical comparison was conducted by one-way ANOVA with a Bonferroni correction. The p value comparisons are DMSO vs. VPA Dox p=0.06, DMSO vs. VPA p= 0.00003, and VPA vs. VPA Dox p= 0.04. **I.** Nuclear height of SAF-A-mAID-mCherry RPE-1 cells with dox induced expression of SAF-A-GFP treated with 0 or 1 uM of dox along with VPA for 24 hours. Data from two biological replicates was graphed as a violin plot. Statistical comparison was conducted by one-way ANOVA with a Bonferroni correction. The p value comparisons are DMSO vs. VPA p=0.002 and DMSO vs. VPA Dox p= 0.002.

### SAF-A is a concentration dependent structural scaffold

If SAF-A acts as a structural scaffold at sites of active transcription, then loss of SAF-A in cells with higher transcriptional activity would be predicted to exhibit an increase in nuclear abnormalities. Previous studies demonstrated that treatment of cells with a global histone deacetylase inhibitor (valproic acid, VPA) causes an increase in mRNA synthesis and weakening of nuclear rigidity^3,16,17^. RPE-1 cells treated with VPA for 24 hours had increased H3K27ac measured by quantitative immunofluorescence and a ∼1.5 µm decrease in nuclear height (Figure S6AB). In addition, EU labeling of 24-hour VPA treated cells showed ∼1.8X increase in nascent transcription compared to untreated control cells (Figure S6C). These results suggest VPA treatment increases the global euchromatin to heterochromatin ratio, resulting in higher transcriptional activity and nuclear softening. To determine if an increase in transcription exacerbates nuclear morphology defects in SAF-A Dep cells, we co-treated SAF-A-mAID-mCherry RPE-1 cells with VPA, doxycycline, and IAA for 24 hours and monitored nuclear morphology. As previously reported, ∼10% of cells treated with VPA had nuclear abnormalities, typically presenting as nuclear blebs (Fig 3C-D). SAF-A depleted cells treated with VPA had a dramatic increase in nuclear abnormalities with ∼20% of the population having shape defects (Fig 3C-D). Qualitatively, nuclear defects in VPA treated SAF-A depletion cells were often severe with multiple nuclear blebs (Fig 3C). This result suggests that cells with higher transcriptional activity are more vulnerable to nuclear shape defects upon SAF-A depletion, supporting a model in which SAF-A functions as a dynamic structural scaffold in conjunction with nascent RNA transcripts.

To determine if increasing SAF-A expression protects against nuclear defects forming in response to elevated transcriptional activity we treated SAF-A-mAID-mCherry cells containing an inducible SAF-A (SAF-A-WT-GFP) rescue construct with doxycycline for 24 hours, resulting in a ∼2-fold increase in SAF-A protein level (Fig 3. F,G). Overexpression of SAF-A reduced VPA induced nuclear abnormalities by ∼60% (Fig 3. H). Nuclear height of VPA treated cells overexpressing SAF-A remained ∼1.5 µM shorter than control cells with no statistical difference compared to VPA only treated RPE-1 cells (Fig 3. I). Therefore, partial rescue of nuclear morphology by SAF-A overexpression in VPA treated cells is independent of nuclear height. Taken together, these results suggest SAF-A is a concentration-dependent structural scaffold at sites of active transcription.

To further evaluate the dose-dependent requirement of SAF-A to preserve normal nuclear morphology we generated a heterozygous SAF-A-AID cell line. One SAF-A allele was modified at the 3′ end using CRISPR-Cas9 to introduce a minimal AID (mini-AID) degron and a SNAP tag into RPE-1 cells constitutively expressing AID2 (TIR1-F74G) (Fig. 4 A,B)^46^. Treatment with 1 µM 5-phenyl-indole-3-acetic acid (5-Ph-IAA) for 24 hours lead to acute depletion of the SAF-A-SNAP-mAID allele resulting in ∼50% reduction in SAF-A protein expression (Fig. 4 C,D). Heterozygous depletion of SAF-A for 24, 48, and 72 hours showed normal nuclear morphology at all timepoints (Fig. 4 E). This result suggests heterozygous depletion of SAF-A in normally cycling cells does not lead to aberrant nuclear shape.

**Figure 4.**
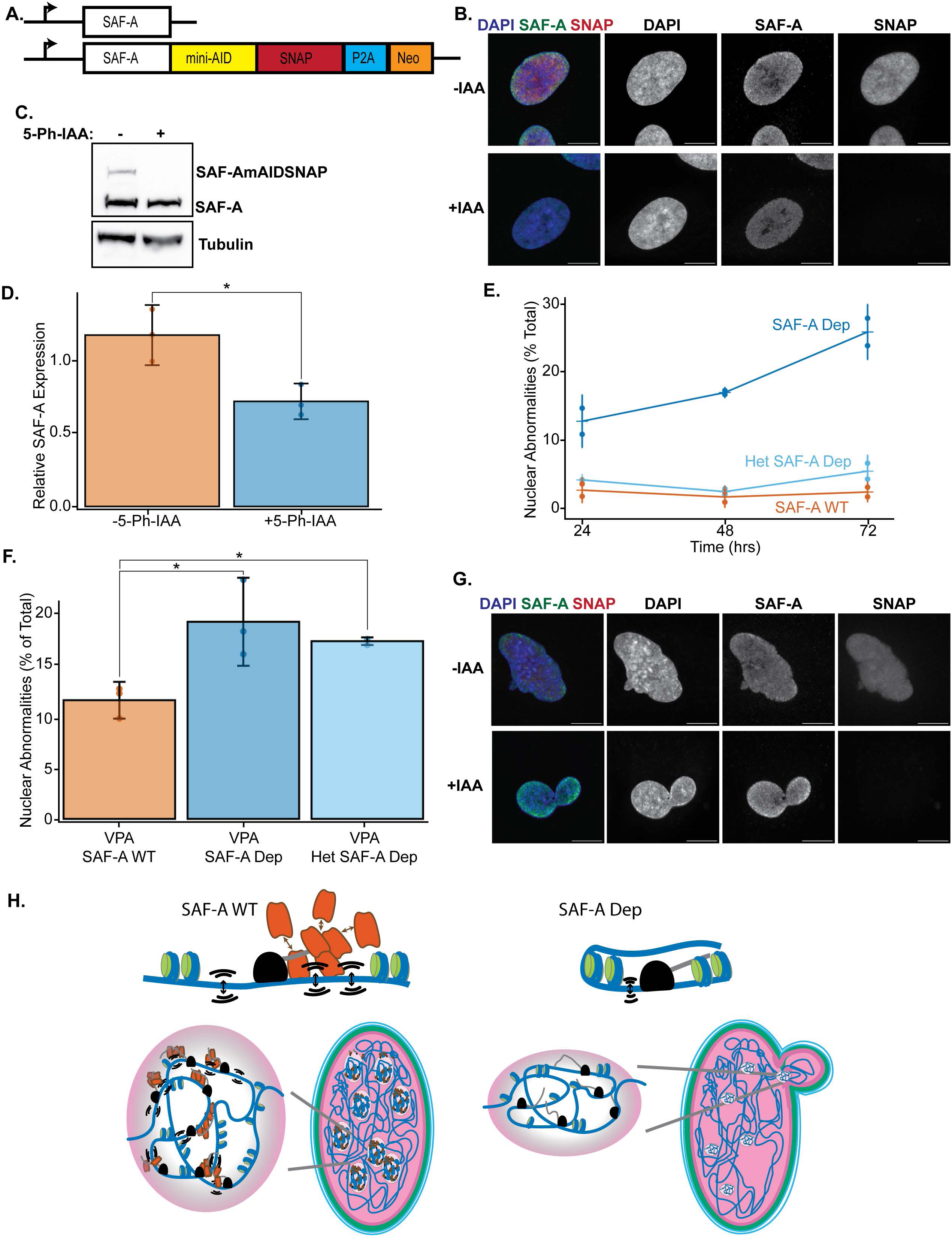
**A.** Schematic of a one copy of endogenous SAF-A tagged with SNAP and mAID in cells containing constitutive TiR1F74G. **B.** Immunofluorescence for SAF-A in heterozygous SAF-A-SNAP-aid cells labeled with SNAP dye and treated with IAA and dox for 24 hours. Images are rendered as a maximum projection of a 3D stack. Bar, 10 µM. **C.** Western blot for SAF-A and Tubulin using extracts from heterozygous SAF-A-mAID-SNAP cells treated with IAA and dox for 24 hours. **D.** SAF-A protein expression normalized to tubulin loading control of western blot analysis shown in A. Data from three biological replicates are shown as a bar graph. Statistical comparison was conducted by an unpaired Student’s t-test with a resulting p value of 0.03. **E.** Quantitation of nuclear abnormalities in 24-, 48-, and 72-hr IAA dox treated heterozygous SAF-A-mAID-SNAP cells (Het SAF-A Dep) and homozygous SAF-A-mAID-mCherry cells (SAF-A Dep). Data from two biological replicates was graphed as percent of cell population with nuclear abnormalities. Statistical comparison was conducted by one-way ANOVA with a Bonferroni correction. The p value comparisons are SAF-A WT vs. Dep 24 hr p= 0.02, 48 hr p= 0.007, 72 hr p= 0.004, SAF-A WT vs. het Dep 24 hr p= 0.7, 48 hr p=0.6, 72 hr p= 0.4. **F.** Quantitation of nuclear abnormalities in IAA dox treated heterozygous SAF-A-mAID-SNAP cells (Het SAF-A Dep) and homozygous SAF-A-mAID-mCherry cells (SAF-A Dep) co treated with VPA for 24 hours. Data from three biological replicates was graphed as percent of cell population with nuclear abnormalities. Statistical comparison was conducted by one-way ANOVA with a Bonferroni correction. The p value comparisons are SAF-A WT vs. Dep p= 0.02 and SAF-A WT vs. het p= 0.05. **G.** Immunofluorescence for SAF-A in heterozygous SAF-A-mAID-SNAP cells labeled with SNAP dye and treated with IAA, dox, and VPA for 24 hours. Images are rendered as a maximum projection of a 3D stack. Bar, 10 µM. **H.** Model of SAF-A as dynamic scaffold that promotes chromatin motion at sites of active transcription.

However, we hypothesized that heterozygous depletion of SAF-A could reduce nuclear mechanical buffering capacity in response to elevated transcription. To test the role of reduced SAF-A concentration in response to elevated transcription we treated SAF-A heterozygous cells with VPA to increase global transcriptional activity for 24 hours. In cells with elevated transcription, heterozygous SAF-A loss led to a dramatic increase in nuclear abnormalities that phenocopied homozygous protein depletion (Fig. 4 F,G). This result suggests that heterozygous loss of SAF-A reduces nuclear mechanical resistance to increased transcriptional activity.

## Discussion

Abnormalities in nuclear shape are caused by an imbalance of tension at the nuclear envelope and are phenotypes observed in cancer and age-related disorders^47,48^. Abnormally shaped nuclei commonly exhibit a global increase in euchromatin with active RNAPII present in nuclear blebs^16,49–51^. RNAPII transcription has been proposed to regulate nuclear mechanical tension, yet the underlying molecular mechanism remains unknown.

Our results demonstrate that SAF-A is a key protein linking RNAPII transcription to increased nuclear rigidity and is required to protect nuclear envelope integrity. Here, we demonstrate that acute depletion of SAF-A causes loss of chromatin motion, formation of nuclear blebs, and compromised nuclear envelope. We identify a dose-dependent requirement for SAF-A to maintain euchromatic architecture under conditions of elevated transcription and nuclear softening. Decreasing global transcription rescued the effects of SAF-A depletion while heterozygous SAF-A loss was highly sensitive to increased transcription. Importantly, a SAF-A mutant that is unable to bind RNA also leads to nuclear morphology defects. Collectively, these results suggest that SAF-A levels must be balanced with the levels of transcription to protect nuclear envelope integrity.

SAF-A is an extremely abundant nuclear protein with estimates ranging from 1-5µM nuclear concentration^52,53^, consistent with a role as a nuclear structural protein. SAF-A is a core component of the ‘nuclear matrix’, a protein-rich insoluble network resistant to high salt or detergent extraction. Nuclear matrix proteins contain nucleic acid binding motifs, with several studies suggesting they are required to maintain nuclear shape, size, and cellular fitness^24,54,55^. The nuclear matrix was initially proposed to be a static structural scaffold; however, FRAP assays clearly demonstrate associated proteins are highly dynamic^28,29,56–59^. Here, we show that association of SAF-A with nascent transcripts forms a dynamic, dose-dependent structural scaffold that modulates chromatin compaction and motion of transcriptionally active regions. Further work is required to identify if additional nuclear matrix proteins such as matrins and SAF-B work cooperatively or independently of SAF-A:RNA scaffolds.

Our findings highlight a structural role of nascent transcripts in preserving the mechanical integrity of the nucleus. We hypothesize that global transcriptional activity generates a dense mesh of SAF-A:RNA interactions to maintain nuclear rigidity. Our single particle tracking of nucleosomes suggest chromatin mobility must be maintained within a homeostatic range. At actively transcribed loci, loading of RNAPII and associated cofactors constrains chromatin mobility^15,43^. This restriction is counterbalanced by dynamic interactions of SAF-A with chromatin and nascent transcripts, which are required to maintain a basal level of chromatin movement.

We propose that dynamic interactions of SAF-A and nascent RNAs maintain dynamic transcriptional hubs, thereby creating a structural meshwork linking euchromatic regions. In this model, SAF-A localizes to sites of active transcription and binds nascent transcripts through its C-terminal RGG repeat domain. RNA binding stimulates oligomerization of SAF-A to form a large complex with RNA and other associated proteins^60^. Multimerized SAF-A complexes promote the opening of euchromatic regions at sites of active transcription which is needed for remodeling and processing of nascent transcripts^27,61^. Loss of SAF-A causes a compaction of euchromatin, yet maintains normal levels of transcription, which results in nuclear envelope stress and loss of nuclear compartmentalization^26^. These results uncover a novel role of RNA–SAF-A networks as regulators of euchromatic architecture and chromatin dynamics to maintain nuclear envelope mechanical tension

### Limitations of this study

While our study identifies SAF-A as a regulator of nuclear morphology and chromatin dynamics in RPE1 cells, further work will be required to validate these results in additional cell types. It is possible that the mechanistic role of SAF-A may differ in cell lines derived from cancerous tissue or expressing different levels of SAF-A. We demonstrated a gradual increase in nuclear abnormalities over 72 hours of SAF-A depletion, however, longer depletion timepoints may provide insight into the essentiality of SAF-A.

## Resource availability

## Lead Contact

Further information and requests for resources and reagents should be directed to and will be fulfilled by the lead contact, Michael D. Blower.

## Materials availability

Plasmids and cell lines generated in this study will be available upon request.

## Data and code availability

All raw reads and summary files are available in GEO under the accession numbers: GSE277216, GSE277217, GSE277219, GSE277221.

## Acknowledgements

The authors thank members of the Blower Lab and Dr. Radhika Subramanian for helpful suggestions. We would like to thank Dr. Matthew Layne and Dr. Cheyanne Frosti for their guidance in preparing polyacrylamide hydrogels. We thank L. Lavis for providing Janelia Fluor dye JFX_650_. We acknowledge that much of the computational work presented in this paper was performed on the Shared Computing Cluster which is administered by Boston University’s Research Computing Services. This work was supported by grants from NIH to M.B. (R01GM122893, R01GM144352) and to A.S.H (R01EB035127, R01CA300848). M.M. is supported by an American-Italian Cancer Foundation Post-Doctoral Research Fellowship and an American Cancer Society Postdoctoral Fellowship (PF-25-1419989-01-PFMBB).

## Author contributions

Conceptualization, K.A and M.D.B.; Methodology, K.A and M.D.B.; Investigation, K.A., M.M., M.D.B.; Visualization, K.A., M.M., M.D.B.; Data curation; M.D.B.; Formal analysis; K.A., M.M., A.S.H., M.D.B.; Funding acquisition M.D.B, A.S.H.; Supervision; M.D.B; Writing; K.A., M.M., A.S.H., M.D.B.

## Declaration of generative AI and AI-assisted technologies in the writing process

During the preparation of this work, the authors used Claude in order to improve clarity of the text. After using this tool, the authors reviewed and edited the content as needed and take full responsibility for the content of the published article.

## Supplemental information

Document S1: Figure S1-S6

## Methods

### Cell culture and drug treatment

hTERT immortalized RPE-1 cells (gift from Brian Chadwick, Florida State University; ATCC CRL-4000) were cultured in DMEM/F12 (Sigma) + 10% FBS (Cytiva) + Pen/Strep. For all auxin-mediated SAF-A homozygous depletion experiments, cells were treated with 1 µM doxycycline (dox, D344, MilliporeSigma) and 1 µM 3-indole acetic acid (IAA, I-5148, MilliporeSigma) for the indicated time period. For auxin-mediated SAF-A heterozygous depletion, cells were treated with 1 µM doxycycline (dox, D344, MilliporeSigma) and 1 µM (5-phenyl-indole-3-acetic acid (5-Ph-IAA, IA1388, MilliporeSigma) for indicated time periods. To induce SAF-A allele exchange, cells were incubated with 1 µM dox and 1 µM IAA for 24 hours. SAF-A overexpression was achieved by incubating cell containing a full length SAF-A rescue construct in complete medium with 1 µM dox for 24 hours. Expression of the NLS-HaloTag reporter construct was induced by incubating cells in complete medium with 1 µM dox for 24 hours. Cells were then incubated with HaloTag Oregon Green Ligand (G2801, Promega) at 0.2 μM for 15 minutes, washed 5 times in complete medium, and incubated in complete medium supplemented with 1 µM dox for 1 hour before live imaging. For serum starvation assays, cells were incubated in serum-free medium for 48 hours. EU labeling cell culture media was achieved by supplementing complete medium with 1 mM of 5-Ethynyl Uridine (50-210-8028, Fisher) for 1 hour prior to fixation. Transcription inhibition was accomplished by incubating cells in medium containing 1 µM triptolide (insert company) or 1 µM LDC000067 (insert company) overnight. For myosin II inhibition, cells cultured in complete medium supplements with 50 µM Blebbistatin (50-226-1421, Fisher) for 24 hours. Inhibition of histone deacetylases was achieved by culturing cells in complete medium supplemented with 2 mM valproic acid (P4543, MilliporeSigma) for 24 hours. mTORC2 was inhibited by culturing cells in complete medium supplemented with 200 nM INK-128 (HY-13328; MedChemExpress) or 5 µM JR-AB2-011 (HY-122022; MedChemExpress) for 24 hours. SNAP labeling was completed by incubating cells in complete medium supplemented with 1 µM of SNAP-Cell® 647-SiR dye (NEB, S9102S) for 30 minutes followed by 5 washes with complete medium and an additional 1-hour incubation prior to fixation.

### Plasmid construction

All plasmids constructed for this publication were created using Infusion cloning. GFP-tagged SAF-A alleles were cloned into the lentiviral expression vector pLVX-TetOne-puro (Takara Bio). The construction of lentiviral plasmids pMB1103 (SAF-Awt-GFP), pMB1109 (SAF-AAA-GFP), pMB1311 (SAF-A-ADD-GFP), pMB1244 (SAF-AΔRGGGFP), and pMB1316 (SAF-AΔSAPGFP) have been previously described^30^. Endogenous targeting of the C terminal SAF-A locus has been previously described. For the SNAP tagged SAF-A degron line, pMB1116 (SAF-A-mCherry-aid) was modified to contain SNAP in place of mCherry. H2B-YFP lentiviral expression vector was ordered from Addgene (#26000). NLS-HaloTag lentiviral expression vector was cloned into a modified pLVX-TetOne vector containing Blasticidin resistance. All plasmid sequences were confirmed by whole plasmid sequencing. Plasmids for PiggyBac-mediated integration of H2B-HALO are previously described ^43^.

### Cell lines

Construction of the SAF-A-mCherry degron line in RPE-1 cells has been previously reported ^30^. The SAF-A-SNAP degron line was built in RPE1 cells expressing TIR1-F74G at the AAVS1 locus. TIR1-F74G was incorporated by nucleofecting 2 μg of pMB1398 (EF1-a promoter-TIR1-F74G-P2A261 Blasticidin resistant gene) and pMB1422 (Cas9-D10A targeting AAVS locus) into 10^6^ cells. Cells recovered for 7 h, followed by addition of pfithrin-α (30 μM; Komarov et al., 1999), pfithrin-μ (10 μM), or no drug for 3 d. Then, cells were selected for by incubating in complete medium containing with 10 μg/ml blasticidin S (Thermo Fisher Scientific). The SAF-A-SNAP degron line was then nucleofected into RPE1 TIR1-F74G cells and selected for by neomycin drug selection. Clones were validated by western blot. Rescue cell lines expressing SAF-A-WT-GFP, SAF-A-ΔSAP-GFP, SAF-A-ΔRGG-GFP, SAF-A-K510A–GFP, or SAF-A-D580A–GFP in SAFA degron RPE-1 background have been previously described. Lentiviruses with YFP-H2B and NLS-HaloTag were synthesized and used for transduction into SAF-A-mCherry degron RPE-1 cells. For drug selection, the transfected RPE-1 cells were selected in medium containing 10 µg/ml of Blasticidin S HCL for the NLS-HaloTag construct only. Individual clones were selected and validated by fluorescent detect of YFP or HaloTag dye. The H2B-HALO SAF-A-mCherry degron line was generated by PiggyBac-mediated genomic integration. SAF-A-mCherry degron cells were transfected with plasmids pASH41^62^ (encoding H2B-HALO) and pASH39 (encoding the PiggyBac integrase). Using using Lipofectamine 3000 (Thermo Fisher, L3000008) according to the manufacturer’s protocol. Following transfection, cells were validated for H2B-Halo expression by flow cytometry.

### Polyacrylamide hydrogel preparation

Polyacrylamide hydrogels of individual discrete stiffness prepared to fit 12 well plates based on a previously published protocol ^37^. Briefly, a 5:4 ratio of 40% acrylamide and 2% bis acrylamide were prepared to make hydrogels with Young’s modulus of 70 kPa, respectively. To ensure gels were cast with even thickness a drop of known volume of the acrylamide solution was pipetted on hydrophobic glass plates pretreated with SUfraSil and flattened by a coverslip during the polymerization process Following polymerization, the gels were derivatized using a dopamine crosslinker (2 mg/ml dopamine 50 mM HEPES buffer pH 8.5) as previously described ^38^. Monomeric collagen (PureCol, Advanced Biomatrix #5005-B) diluted in sterile 1X PBS at 0.05 mg/ml was delivered to each well and incubated overnight at 4°C.

### Cell fixation and Immunofluorescence

Cells were fixed with 4% PFA in 1× PBS for 10 min at room temperature, then permeabilized in PBS + 0.5% Triton X-100 for 15 min at room temperature. For PCNA immunofluorescence cells were permeabilized in CSKT for 1 minute prior to fixation in 4% PFA in PBA for 10 minutes. Cells were incubated in blocking buffer (PBS + 1% ultrapure BSA + 2% v/v Tween) with primary antibody for 1 hour at 37°C or overnight at 4°C. After 3 × 5 min washes with PBS cells were incubated with secondary antibody in blocking buffer for 1 hour at 37°C. Cells were then washed 3 × 5 min with PBS-T. For EU detection, the previously published protocol was followed ^30^.

The following primary antibodies were used in this study for immunofluorescence: rabbit anti LaminB1 (12987, ProteinTech), mouse anti Mitosin (610768, BD Biosciences), mouse anti H3K27ac (AB23516, Invitrogen), mouse Ki67 (61098, BD Biosciences), rabbit p21 (10355-1-AP, ProteinTech), rabbit anti RNAPII Ser2P (5095, Abcam), rabbit anti RNAPII Ser5P (5131, Abcam), rabbit anti H3K9me3 (39062, Active Motif).

Secondary antibodies used in immunofluorescence assays were as follows: donkey anti-rabbit Alexa Fluor 488 (711-545-152, Jackson Immunoresearch), donkey anti-rabbit Alexa Fluor 647 (709-605-149, Jackson Immunoresearch), donkey anti-mouse Cy3 (715-165-150, Jackson Immunoresearch), and donkey anti-mouse Alexa 647 (715-605-150, Jackson Immunoresearch).

### Immunoblotting

RPE1 cells were trypsinized and harvested, washed with PBS, and suspended in INSERT EMILY’s Buffer and lysed by sonication. Protein concentration of extracts was measured by Bradford assay. Equal volume of protein concentration for sample conditions were mixed into 1x Laemmli sample buffer and heating for 5 min at 95°C. Proteins were separated on a 8% SDS–PAGE gel, handmade and transferred to Immobilon (Millipore) by Trans-Blot Turbo (BIORAD) with 2.5 A for 12 min. Membranes were washed with PBS + 2 vol/vol% Tween 20 and blocked with PBS + 2 vol/vol% Tween 20 supplemented with 5% skim milk for 1 hour at room temperature. Then, primary antibody was supplemented to the blocking solution, and membranes were incubated overnight at 4°C, shaking. Then, membranes were washed with PBS + 2 vol/vol% Tween 20 for 5 min three times and incubated with secondary antibody in PBS + 2 vol/vol% Tween 20 and blocked with PBS + 2 vol/vol% Tween 20 supplemented with 5% skim milk for 1 h at room temperature. Then, membranes were washed with PBS + 2 vol/vol% Tween 20 for 5 min three times and incubated with Immobilon ECL Ultra Western horse radish peroxidase (HRP) Substrate (Millipore) for 5 min, then scanned in a Chemidoc MP imager (Bio-Rad). For the primary antibodies, 1/X of anti-SAFA from mouse (Santa Cruz, X) and 1/10,000 of anti–α-Tubulin from mouse (Sigma, 9026). For secondary antibody, HRP-conjugated antibodies were used.

### Image acquisition and analysis

Images were acquired using a Nikon A1R confocal microscope with a 60 x 1.4NA Vc lens. For live cell analysis, images were captured using a scan zoom of 2x (140 nm/pixel) using a resonant scanner with Z stacks spaced 5 µm apart. Images were acquired every 5 minutes.

For live cell analysis, nuclear circularity was quantified by a script in Fiji by thresholding maximum projected images to identify NLS-HALO positive regions and calculating the circularity using the Shape Descriptors feature. Nuclei in live cell image analyses were identified by a script in Fiji which thresholds maximum projected images and measures the fluoresence intensity and circularity of NLS-HALO positive segmented regions. Cytoplasmic NLS-HALO fluorescence intensity was measured by subtracting nuclear signal from the total cell fluorescence intensity in Fiji. For fixed analysis, images were captures using a scan zoom of 2x or 4x (140 or 70 nm/pixel) using a Galvano or Resonant scanner. Images were acquired as Z stacks spaced 0.2 µm apart. All experiments were repeated for a minimum of two biological replicates. The number of biological replicates and total number of analyzed cells are included in each figure legend. Nuclear fluorescence intensity was measured using a script in Fiji with nuclei defined by DAPI-positive regions. For each experiment, nuclear fluorescence intensity was normalized to the average of untreated control cells. Nuclear height was calculated by constructing x-y orthogonal view projections in Fiji and measuring the distance from the top to bottom of

Dapi stained nuclei. For each cell, the average of three nuclear height measurements was the recorded value used for statistical analysis. Data are shown as mean ± standard error of biological replicates. For multiple comparisons, a one-way ANOVA with a post-hoc Tukey test was conducted. For single comparisons, a two-tailed unpaired or paired t-test of mean intensities was performed, as indicated in figure legends.

### Single Particle Tracking (SPT) Cell treatment and labeling

For chromatin motion measurements, we investigated RPE-1 cells under four conditions: untreated, SAF-A depletion (SAF-A KD), transcription inhibition with triptolide (TRP), and SAF-A KD combined with TRP. For SAF-A KD, OsTIR1 expression was first induced by adding doxycycline (Dox, 1 µg/mL) at 48 h before single-particle tracking (SPT), and next SAF-A depletion was induced by adding auxin (IAA, 500 µM) at 24 h before SPT. For transcription inhibition, TRP (1 µM) was added 24 h before SPT. For the combined SAF-A KD + TRP condition, Dox, IAA, and TRP were added following the same timing and doses of the individual treatments. All treatments were maintained throughout the SPT experiment.

To track histone H2B-Halo mobility, we seeded cells into 24-well glass multi-well plates (Cellvis, P24-1.5H-N) three days before imaging, at 30% confluency. One day before imaging cells were incubated for 30 min at 37 °C with cell medium supplemented with the Halo ligand JFX_650_ ^63^at low concentration (0.75 pM) allowing for sparse labeling and therefore for visualization of H2B-Halo single molecules. After labeling, cells were washed three times in PBS and grow in regular medium. To facilitate nuclei localization, on the day of imaging cells were incubated for 15 min at 37 °C with cell medium supplemented with Hoechst 33342 (0.5 µM; Santa Cruz Biotechnology, sc-495790), followed by three PBS washes. Cells were next imaged in phenol-red-free DMEM/F-12 (Thermo Fisher Scientific, 21041025) supplemented with supplemented with 10% fetal bovine serum (Avantor Seradigm, 89510-186, lot 190B20), 100 µg/mL penicillin-streptomycin (Thermo Fisher, 15140122) and 2 mM GlutaMAX (Thermo Fisher, 35050061).

### SPT acquisitions

H2B-Halo tracking was performed largely as previously described^43^. SPT was performed on a custom-built^64^ Nikon Ti2-E inverted microscope equipped with a 100x/1.49 NA oil-immersion apochromatic objective and Prime 95B sCMOS cameras (Teledyne Photometrics; 110 nm effective pixel size). To excite individual H2B-Halo molecules, we applied highly inclined and laminated optical sheet (HILO) illumination^65^, with an illumination angle set to 57° using an iLas 2 motorized dual-galvo system (Gataca Systems). To maintain physiological imaging conditions, samples were kept at 37 °C, 5% CO_2_, and controlled humidity using an Okolab stage-top chamber.

Imaging was performed using a 640-nm Coherent Genesis excitation laser at 10% laser power and an acousto-optic tunable filter (AOTF) set to 100%, on a field of view of 512 x 512 pixels which enables multiple cells to be imaged at once. HILO was performed in ellipse mode, with the excitation beam tracing an elliptical scan (6.67 ms per rotation) to achieve uniform illumination of the field of view.

To track chromatin dynamics over 3 orders of magnitude in time, two acquisition regimes were used as previously described^43^ to overcome photobleaching: a fast regime with a 100-ms interval between consecutive frames (10 frames/s) and a slow regime with a 2-s interval (0.5 frames/s), with each movie being 450 frame-long. To reduce contributions from rapidly diffusing, non-chromatin-bound molecules, the 640-nm excitation laser was pulsed for 86.71 ms per frame, corresponding to 13 cycles of the HILO ellipse scan.

Two snapshots of Hoechst-labeled nuclei (405-nm excitation, Genesis 5%, AOTF 100%) were taken, one immediately before and one immediately after each SPT movie, to provide reference images of nuclei. A single snapshot of SAF-A (561-nm excitation, Genesis 5%, AOTF 100%) was taken immediately before each SPT movie to monitor SAF-A expression levels.

### SPT preprocessing

Since a single SPT field of view accommodates multiple cells, a region of interest was manually drawn around each nucleus to discriminate different cells. Next, to identify individual H2B-Halo molecules, SPT movies were analyzed in ImageJ using TrackMate (v7.14.0)^66^. Single particles were detected in each frame using a Laplacian-of-Gaussian filter with a maximum particle diameter of 0.8 µm and a minimum quality score of 3, reflecting spot brightness how closely the spot size matches the specified diameter. To link corresponding detections into trajectories, the LAP algorithm^67^ was used with maximum allowed frame-to-frame displacements of 300 nm (for 100-ms acquisitions) or 450 nm (for 2-s acquisitions), chosen as previously described^43^. TrackMate trajectories were exported as XML files containing the x-y coordinates and frame number of each localization.

### Chromatin dynamics analysis

To quantify chromatin dynamics, H2B-Halo trajectory processing and mean squared displacement (MSD) analysis were performed using our previously described chromatin dynamics analysis pipeline^68^ available on GitHub: https://github.com/ahansenlab/chromatin_dynamics.

Briefly, adjacent H2B-Halo trajectories *x*_1_(*t*) and *x*_2_(*t*) from the same nucleus were paired to calculate the 2-point MSD, which measures their relative movement while correcting for cell motion:

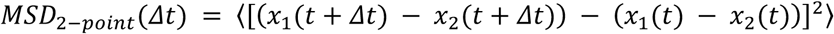

Only trajectories pairs with at least 20 shared frames were used, to ensure sufficient temporal overlap. Pairs more than 3 µm apart on average were excluded to limit motion potentially due to nuclear deformation. Finally, redundant trajectory pairs were removed, so that the same particle motion was not counted multiple times.

Next, to quantify how much a single chromatin locus moves as a function of lag time, we plotted 0.5 x the 2-point MSD, which corresponds to the 1-point MSD under the assumption that the two paired loci have independent motion. For each condition and acquisition interval, trajectories from the three biological replicates were pooled before calculating the ensemble MSD, yielding one MSD curve across all measured trajectories.

To quantify how chromatin motion scales with lag time, trajectories from the 100-ms and 2-s acquisitions were jointly fitted using our previously described Bayesian MSD fitting framework^43^. We assumed an underlying power-law MSD:

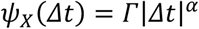

where *Γ* is the MSD prefactor and *α* ∈ [0, 2) is the MSD exponent, with *α* < 1 indicating subdiffusion and smaller values corresponding to stronger subdiffusion. To account for motion blur and localization error, we fitted the expected measured MSD:

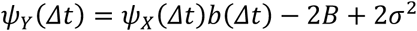

with

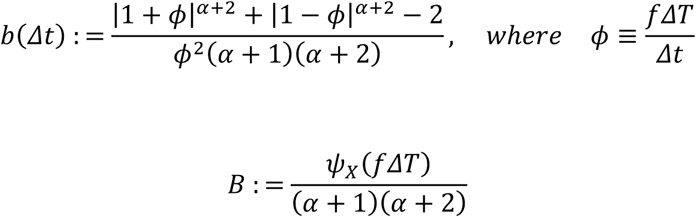

where *fΔT* is the exposure time (86.71 ms), and σ is the localization error. Thus, the fit explicitly accounts for motion blur caused by the finite exposure time and for localization uncertainty.

Given the assumption that chromatin dynamics follow a single power law across the measured time scales, the joint fit used the same *Γ* and *α* for both acquisition regimes. However, to assess whether α was consistent between acquisition regimes, the 100-ms and 2-s datasets were also fitted separately.

Finally, to summarize the effect of each treatment on chromatin motion, we calculated the relative mobility by averaging the logarithm of the treatment-to-control MSD ratio over logarithmic lag time:

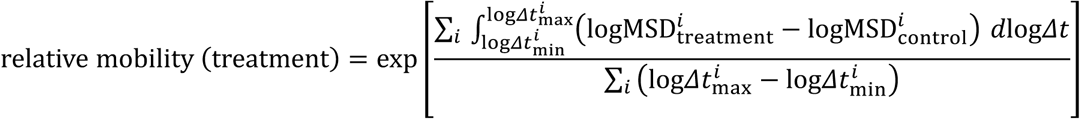

where *i* denotes the SPT acquisition regime (100 ms or 2 s). Exponentiation converts this log-space average into a fold-change relative to control, with values above or below 1 indicating an overall increase or decrease in chromatin mobility, respectively.

### Cell numbers and replicates for each experiment

**Figure 1. B.** Cell n for scoring of nuclear abnormalities is: -IAA 24 hr n=45 and n=69, +IAA 24 hr n=120 and n=80,-IAA 48 hr n=101 and n=53, +IAA 48 hr n=88 and n=97, -IAA 72 hr n=82 and n=55, and +IAA 72 hr n=110 and n=55. **G.** Cell n for nuclear height quantification are SAF-A WT glass n=19, n=18, and n=20 SAF-A Dep glass n=40, n=26, and n=16, SAF-A WT collagen1 n=16 and n=27, SAF-A Dep collagen1 n=33 and n=31, SAF-A WT kPa70 n=28 and n=27, SAF-A Dep kPa70 n=33 and n=23. **I.** Cell n for scoring are SAF-A WT glass n=189 and n=195, SAF-A Dep glass n=165 and n=193, SAF-A WT collagen1 n=170 and n=110, SAF-A Dep collagen1 n=170 and n=110, SAF-A WT kPa70 n=180 and n=114, SAF-A Dep kPa70 n=180 and n=134.

**Figure 2. E.** Cell n for scoring are 10% FBS SAF-A WT n=177 and n=115, 0% FBS SAF-A WT n=112 and n=114, 10% FBS SAF-A Dep n=155 and 114, 0% FBS SAF-A Dep n= 53 and 122. **F.** Cell n for scoring are SAF-A WT n=174, n=245, and n=236, SAF-A Dep n=168, n=235, and n=184, SAF-A Dep LDC n=296, n=206, and n=244, SAF-A Dep TRP n= 243,n=240, and n=199.

**Figure 3. B.** Cell n for scoring are: SAF-A-GFP 24= 154,104, and 142 48= 153, 103, and 108. 72 hr= 157, 101, and 127, SAF-A Dep 24= 237, 219, and 148, 48= 183, 208, and 139, 72= 156, 153, and 149, SAF-A K510A-GFP 24= 153, 102, and 326, 44= 155, 102, and 446, 72= 152, 100, and 296, SAF-A D580A-GFP 24= 152, 100, and 203, 48= 164, 102, and 491, 72= 157, 107, and 482, SAF-A ΔSAP-GFP 24= 126, 207, and 406, 48= 160, 216, and 256, 72= 158, 207, and 293, 24= 98, SAF-A ΔRGG -GFP 89, and 320, 48= 154, 103, and 162, 72= 159, 113, and 201. **D.** Cell n for scoring are SAF-A WT n= 248, 380, and 414, SAF-A Dep n= 231, 321, and 344, SAF-A WT VPA n= 266, 321, and 270, SAF-A Dep VPA n= 144, 345, and 228. **H.** Cell n for scoring are DMSO= 186, 120, 201, 209, 126, and 136, VPA= 142, 158, 123, 143, 152, 136 VPA Dox = 100, 171, 198, 192, 162, and 131. **I.** Cell n for scoring are: DMSO= 34 and 36, VPA= 23 and 36 VPA Dox= 25 and 33.

**Figure 4. E.** Cell n for scoring are: SAF-A WT 24 hr= 250 and 217, 48 hr= 195 and 190, 72 hr= 220 and 230, Het SAF-A Dep 24 hr= 185 and 210, 48 hr= 120 and 182, 72 hr= 192 and 153, SAF-A Dep 24 hr= 237 and 219, 48 hr= 183 and 208, 72 hr= 156 and 153. **F.** Cell n for scoring are: SAF-A WT= 266, 321, and 270, SAF-A Dep= 144, 345, and 228, Het SAF-A Dep= 116, 120, and 128.

**Figure S1. B.** Cell n for quantification is: -IAA 24 hr n=243, n=192, and n=101, +IAA 24 hr n=243, n=199, and n=50, -IAA 48 hr n=188, n=179 and n=58, +IAA 48 hr n=182, n=222, and n=74, -IAA 72 hr n=223, n=179, and n= 59, and +IAA 72 hr n=140, n=150, and n=100. **C.** Cell n for scoring of nuclear abnormalities is: -IAA 24 hr n=250 and n=217, +IAA 24 hr n=237 and n=219,-IAA 48 hr n=195 and n=190, +IAA 48 hr n=183 and n=208, -IAA 72 hr n=220 and n=230, and +IAA 72 hr n=156 and n=153. **G.** Cell n for quantitation is: Glass n= 19, 18, and 20, Blebbistatin n= 39, 14, Collagen n= 16, 27, KPa70 n=28, 27. **H.** Cell n for quantification is WT DAPI n= 99, 61, 40, Dep DAPI n= 93, 73, 36, WT H3K9me3 n= 47, 63, 48, Dep H3k9me3 n= 33, 31, 24, WT RNAPII Ser2p n= 76, 48, Dep RNAPII Ser2p n= 28, 42, WT RNAPII Ser5p n= 78, 51, Dep RNAPII Ser5p n= 28, 43.

**Figure S3. A.** Cell n for quantification is: normal 24 hr n=34, n=31, and n=38, abnormal 24 hr n=39, n=19, and n=27, normal 48 hr n=34, n=28, and n=26, abnormal 48 hr n=57, n=22, and n=25, normal 72 hr n=44, n=45, and n= 32, and abnormal 72 hr n=45, n=32, and n=25. **D.** Cell n for quantification is normal n=36, n=36, and n=33, abnormal n=15, n=15, and n=21. **E.** Cell n for quantification is wildtype rescue: DMSO n= 114, n=140 INK-128 n=107, n=128 JR-AB2 n=105, n=131 and SAF-A depletion: DMSO n=116, n-133, n=119 INK-128 n=253, 78 JR-AB2 n=231, n=55, n=36.

**Figure S5. B.** Cell n for quantitation is: 10% FBS n=104, n=58, and n=93, 0% FBS n=63, n=65, and n= 93. **C.** Cell n for quantitation is: 10% FBS n=135, n=114, and n=50, 0% FBS n=51, n=92, and n= 54. **E.** Cell n for quantitation is: Untreated n= 50, 63, and 59, LDC= 61, 39, and 45, Triptolide= 44, 43, 43.

**Figure S6. A.** Cell n for quantitation is: SAF-A WT n= 19, 18, and 20, SAF-A Dep n= 40, 26, and 16, SAF-A WT VPA n= 21 and 20, SAF-A Dep VPA n= 30 and 22. **B.** Cell n for quantitation is: SAF-A WT n= 36, 41, 59, and 95, SAF-A Dep n= 26, 43, 43, and 52, SAF-A WT VPA n= 39, 48, 74, and 13, SAF-A Dep VPA n= 30, 32, 35, and 36. **C.** Cell n for quantitation is: Untreated n=50 and n=54 VPA n=51 and n= 55.

---

**Figure S1.**
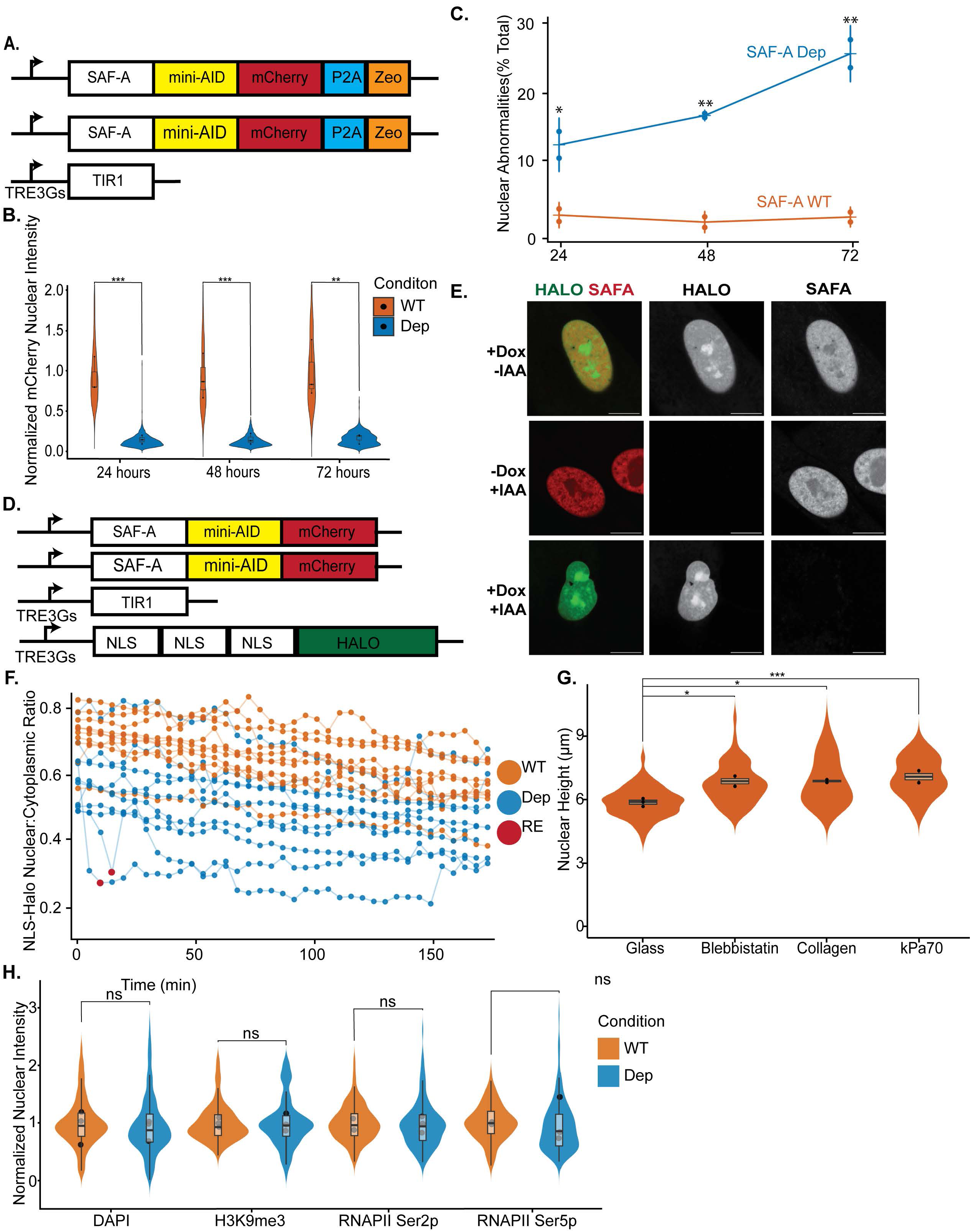
**A.** Schematic of auxin inducible degron system with endogenous SAF-A tagged with mini-AID mCherry in RPE1 cells containing doxycycline inducible Tir1. **B.** Quantification of normalized mCherry fluorescence in 24-hour SAF-A depleted cells. Data from three biological replicates is visualized as a violin plot with the mean of each replicate indicated by a closed circle. Statistical comparison of normalized mCherry fluorescence was conducted using a one-way ANOVA followed by Tukey’s test with a Bonferroni correction. The p value comparisons of -IAA and +IAA were: 24 hr p=0.004, 48 hr p=0.01, 72 hr p= 0.02. **C.** Scoring of nuclear morphology defects of RPE-1 cells treated for 24, 48, and 72 hours with doxycycline and IAA treatment. Image stacks were acquired for at least 150 nuclei per condition for each timepoint for two biological replicates. Mean of the total percent of cells with nuclear abnormalities are plotted as closed circles. Statistical comparison of percent nuclear abnormalities was conducted using a one-way ANOVA followed by Tukey’s test with a Bonferroni correction. The p value comparisons of -IAA and +IAA were: 24 hr p= 0.041, 48 hr p= 0.002, 72 hr p= 0.008. **D.** Schematic of a doxycycline include x3NLS-Halo reporter incorporated into SAF-A-mCherry AID RPE-1 background. **E.** OregonGreen NLS-HALO and mCherry fluorescence of live SAF-A-mAID-mCherry RPE-1 treated with doxycycline and IAA. Images are rendered as maximum projections of a 3D stack. Bar, 10 µm. **F** Quantification of the ratio of nuclear to cytoplasmic NLS-HALO fluorescence intensity (Nuc/Cyto) over a 3-hour live cell imaging time course. Ten cells were monitored for SAF-A wildtype and SAF-A depletion conditions. Nuclear Rupture (RE) events are denoted as a red circle. **G.** Nuclear height comparison of SAF-A-mAID-mCherry cells seeded on glass, collagen, or 70 kPa polyacrylamide hydrogel. Cells seeded on glass were either untreated or treated with blebbistatin. Data from at least two biological replicates are presented as violin plots. Statistical comparison was conducted by one-way ANOVA with a Bonferroni correction. The p value comparisons are Glass vs Blebbistatin p=0.02, Glass vs Collagen p=0.02, Glass vs. kPa70 p= 0.004. **H.** Quantification of nuclear fluorescence intensity of immunofluorescence for H3K9me3. RNAPII Ser2p, RNAPII Ser5p, and DAPI staining in SAF-A-mCherry-Aid cells treated with auxin for 24 hours. Data from three biological replicates is visualized as a violin plot with the mean of each replicate indicated by a closed circle. Statistical comparison of normalized fluorescence for each target was conducted using a unpaired Student’s t-test between WT and Dep groups for each target, resulting in p values of DAPI p= 0.8, H3K9me3 p= 0.9, RNAPII Ser2p p= 1.0, RNAPII Ser5p p=0.9.

**Figure S2.**
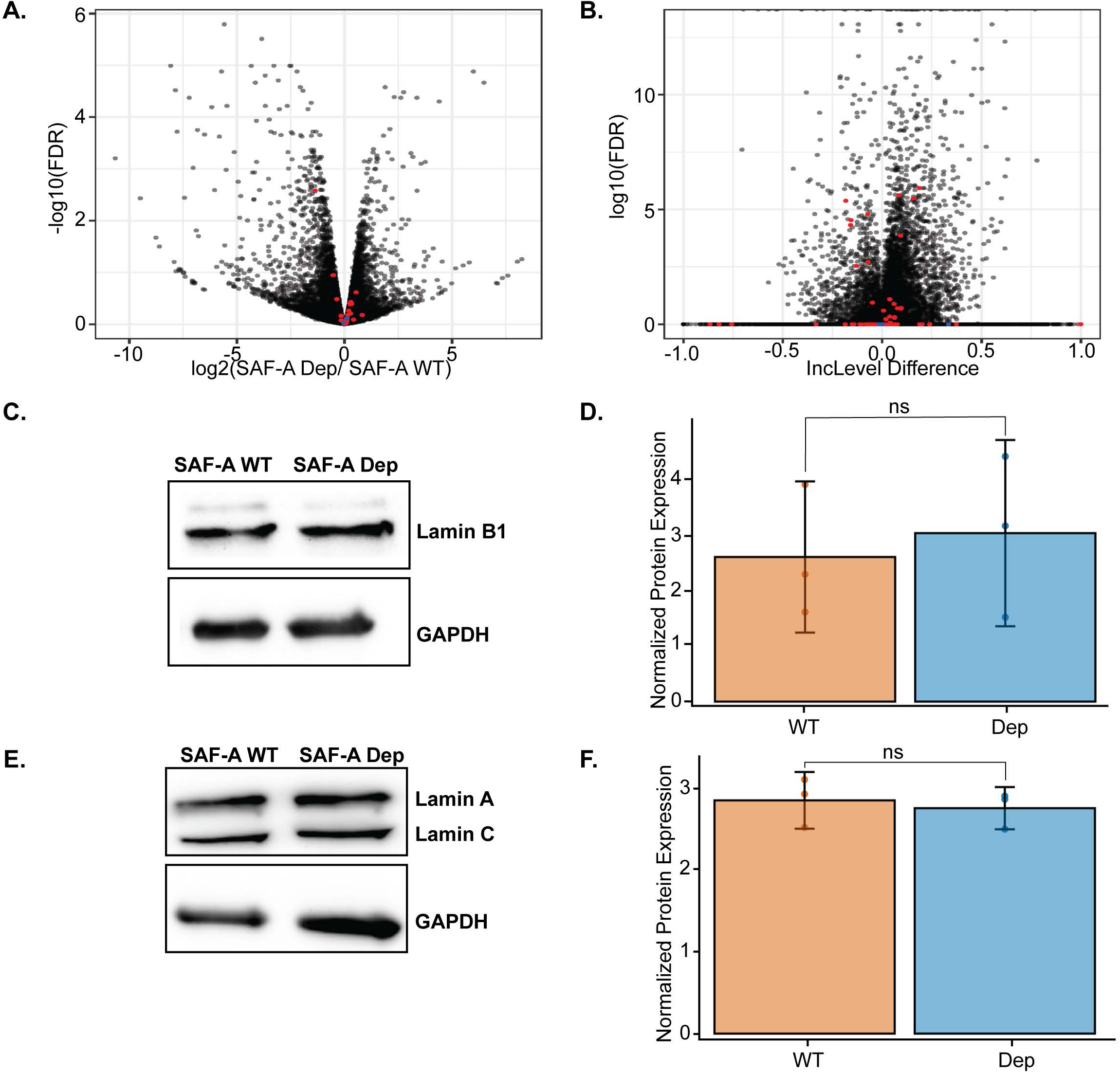
**A.** Gene expression level quantification from RNA-SEQ analysis of SAF-A-mAID-mCherry cells treated with auxin compared to untreated control cells. Lamin proteins are represented as blue circles. Nuclear envelope structural proteins are represented as red circles. **B.** Exon inclusion level differences (RPE1-SAF-A depleted) derived from rMATS analysis of RNA-SEQ analysis of SAF-A-mAID-mCherry cells treated with auxin compared to untreated control cells. Lamin proteins are represented as blue circles. Nuclear envelope structural proteins are represented as red circles. **C.** Western blot analysis of Lamin B1 expression in SAF-A-mAID-mCherry cells treated with or without auxin for 24 hours. GAPDH used as a loading control. **D.** Quantification of western blot displayed in C. Three biological replicated were included, statistical comparison of WT vs. Dep was conducted using an unpaired Student’s t test with a resulting p value of X. **E.** Western blot analysis of Lamin A/C expression in SAF-A-mAID-mCherry cells treated with or without auxin for 24 hours. GAPDH used as a loading control. **F.** Quantification of western blot displayed in E. Three biological replicated were included, statistical comparison of WT vs. Dep was conducted using an unpaired Student’s t test with a resulting p value of 0.7.

**Figure S3.**
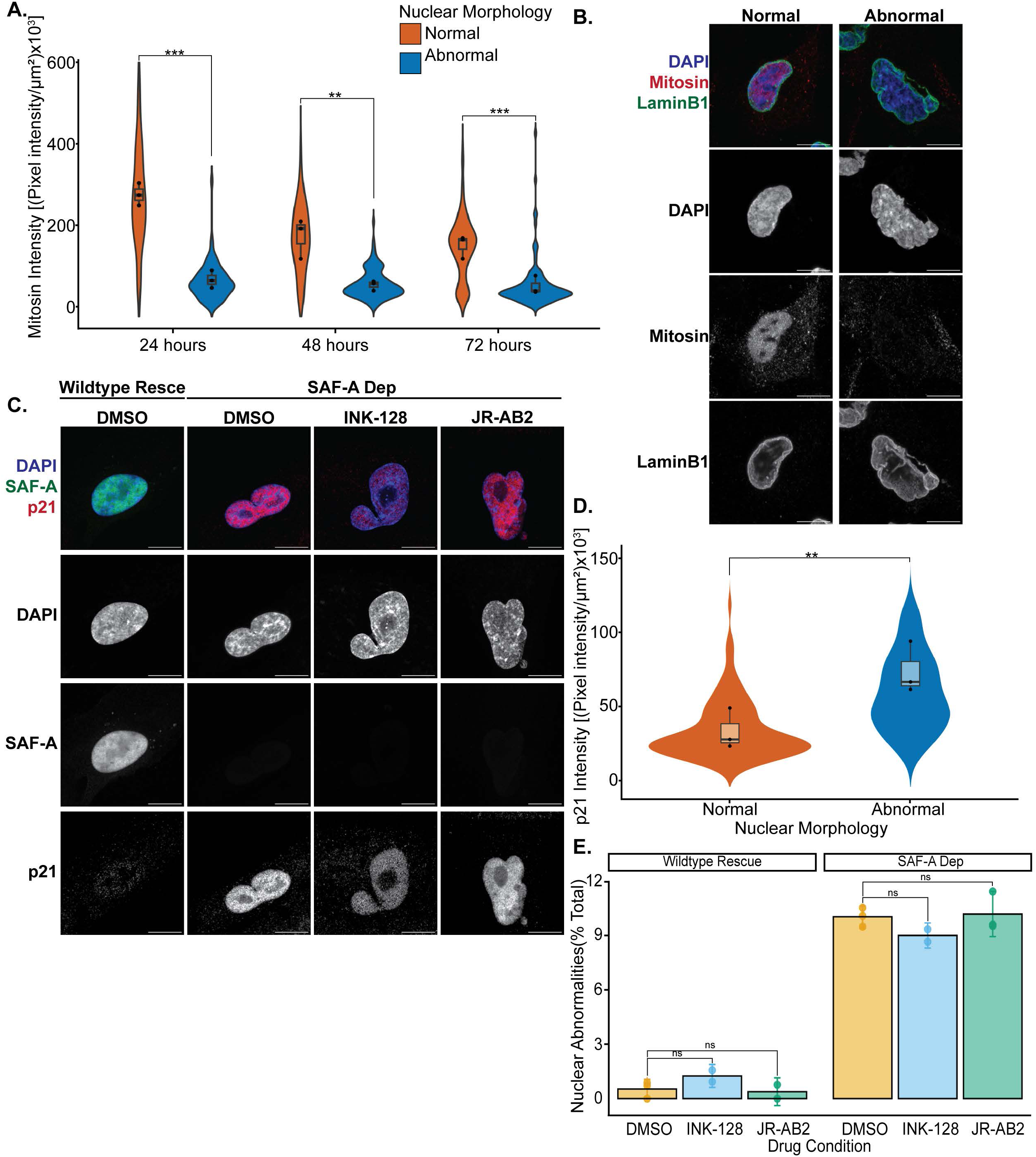
**A.** Quantitation of mitosin fluorescence intensity in 24-, 48-, and 72-hour SAF-A depleted cells. Cells from three biological replicates were stratified based on nuclear abnormalities and compared by running a paired t-test with a Bonferroni correction. The p value comparisons of normal and abnormal were: 24 hr p=0.0005, 48 hr p=0.01, 72 hr p= 0.009. **B.** Imaging of immunofluorescence for Mitosin and laminB1 with DAPI staining in 24-hour SAF-A depletion cells. Images are rendered as maximum projections of a 3D stack. Bar, 10 µm. **C.** Imaging of immunofluorescence for p21 and SAF-A-mCherry fluorescence with DAPI staining in 24-hour SAF-A depleted cells co-treated with INK-128 or JR-AB2. Images are rendered as maximum projections of a 3D stack. Bar, 10 µm. **D.** Quantification of p21 fluorescence intensity in 24-hour SAF-A depleted cells. Data from three biological replicates was stratified by nuclear morphology and compared by conducting a paired t-test with a Bonferroni correction. The p value comparison of normal and abnormal was p= 0.007. **E.** Scoring of nuclear abnormalities as percent total of the population in 24-hr SAF-A depletion or SAF-A wildtype rescue co-treated with DMSO, INK-128, or JR-AB2. Data from at least 2 biological replicates is represented as a bar graph with mean replicate values shown as closed circles. Statistical comparison was conducted by one-way ANOVA with a Bonferroni correction. The p value comparisons are SAF-A WT vs. Dep p= 0.005, SAF-A WT vs. SAF-A WT blebbistatin p= 0.04, SAF-A Dep vs. SAF-A Dep blebbistatin p=0.0002, SAF-A WT blebbistatin vs. SAF-A Dep blebbistatin p=1.0.

**Figure S4.**
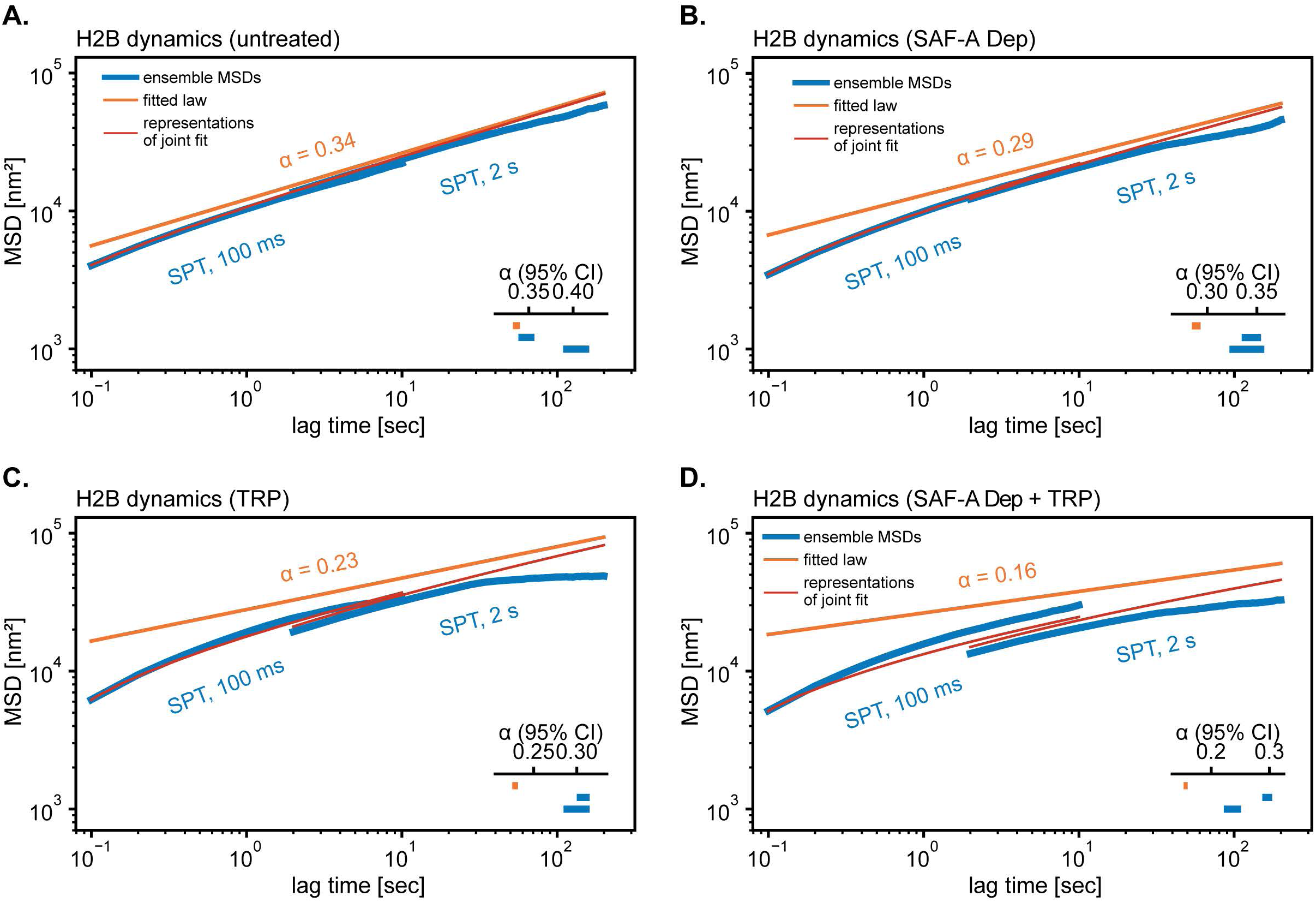
**A-D.** Ensemble mean squared displacements (MSDs) and power-law fits of H2B-Halo in RPE-1 cells across treatment conditions and three biological replicates. H2B-Halo trajectories were acquired at 100-ms and 2-s time lags and combined to span a broader dynamic range for untreated cells (A; 100 ms: n_cells_ = 59, n_tracks_ = 3138; 2 s: n_cells_ = 57, n_tracks_ = 3103), SAF-A knockdown (SAF-A KD, B; 100 ms: n_cells_ = 46, n_tracks_ = 2461; 2 s: n_cells_ = 48 n_tracks_ = 2888), triptolide treatment (TRP, C; 100 ms: n_cells_ = 58, n_tracks_ = 5815; 2 s: n_cells_ = 54, n_tracks = 4267), and combined SAF-A depletion and triptolide treatment (SAF-A KD + TRP, D; 100 ms: n_cells_ = 50, n_tracks = 4091; 2 s: n_cells_ = 49, n_tracks_ = 4042). Ensemble MSDs are shown in blue, the joint power-law fit across both acquisition regimes in orange, and the fits to the individual 100-ms and 2-s datasets in red. Insets show the 95% credible intervals for the exponent α from the joint fit (top, orange), followed by the separate fits to SPT at 100 ms and 2 s. Average mobility relative to untreated control is shown in the inset at the bottom

**Figure S5.**
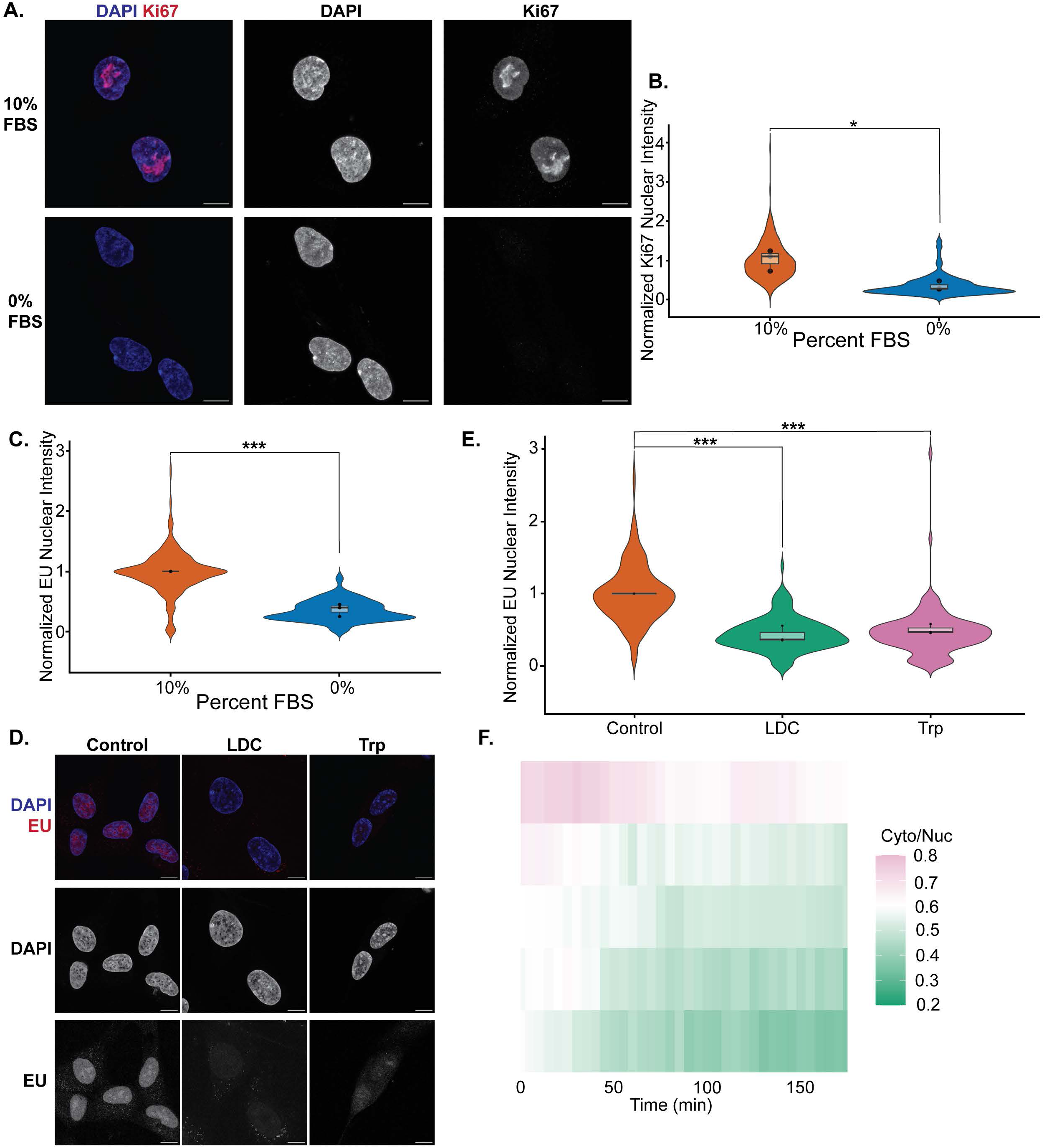
**A.** Immunofluorescence for Ki67 and DAPI staining of RPE1 cells cultured in 10% or 0% FBS for 48 hrs. Images are rendered from maximum projections of a 3D stack. Bar, 10 µM**. B.** Quantitation of Ki67 nuclear fluorescence intensity from experimental conditions in D. Data from three biological replicates are normalized to the average intensity of RPE1 cells cultured in 10% FBS. Statistical comparison was conducted by an unpaired t-test result with p=0.03. **C.** Quantitation of Cy3 nuclear intensity of Cy3-5-EU labeled nascent transcripts in RPE1 cells cultured in 10% or 0% FBS. Data from three biological replicates are normalized to the average intensity of RPE1 cells cultured in 10% FBS. Statistical comparison was conducted by an unpaired t-test result with p=0.009. **D.** DAPI staining of Cy3-5-EU labeled nascent transcripts in RPE1 cells incubated overnight with LDC or Triptolide (Trp). Images are rendered as maximum projections of a 3D stack. Bar, 10 µM. **E.** Quantification of experimental conditions in panel D. Data from three biological replicates are normalized to the average intensity of untreated RPE1 cells. Statistical comparison was conducted by one-way ANOVA with a Bonferroni correction. The p value comparisons are Untreated vs. LDC p= 0.0002 and Untreated vs. Triptolide p=0.005. **F.** Quantitation of nuclear to cytoplasmic ratio of NLS-HALO signal over a 3-hour time course with imaging every 5 minutes. Cells were treated with dox, IAA, and LDC for 24 hours (LDC, n=5).

**Figure S6.**
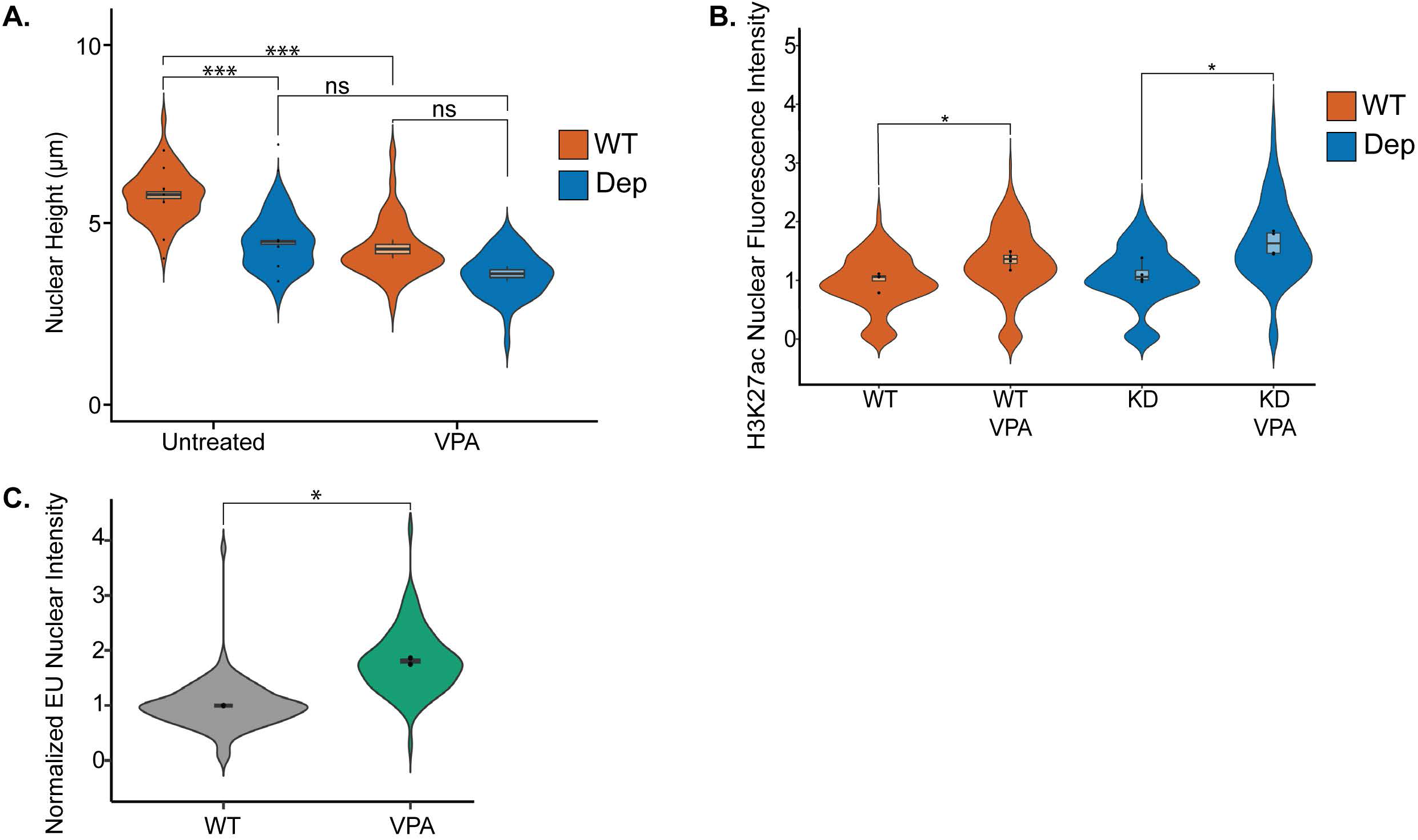
**A.** Nuclear height comparison of SAF-A-mAID-mCherry cells treated with IAA and dox with or without the presence of VPA for 24hrs. Data from at least two biological replicates are presented as violin plots. Statistical comparison was conducted by one-way ANOVA with a Bonferroni correction. The p value comparisons are SAF-A WT vs SAF-A Dep p= 0.005, SAF-A WT vs SAF-A WT VPA p= 0.005, SAF-A Dep vs SAF-A Dep VPA p= 0.09, SAF-A WT VPA vs. SAF-A Dep VPA p= 0.3**. B.** Nuclear intensity of H3K27ac immunofluorescence signal of SAF-A-mAID-mCherry cells treated with IAA and dox with or without the presence of VPA for 24hrs. Data from four biological replicates are normalized to the average intensity of untreated RPE1 cells with SAF-A WT. Statistical comparison was conducted bya unpaired Student’s t-test. The p value comparisons are SAF-A WT vs. SAF-A WT VPA p= 0.01 and SAF-A Dep vs. SAF-A Dep VPA p= 0.01. **C.** Quantitation of nuclear intensity of Cy3-5-EU labeled nascent transcripts in RPE1 cells treated with VPA for 24 hours. Data from two biological replicates are normalized to the average intensity of untreated RPEs. Statistical comparison was conducted by an unpaired t-test result with p=0.05.

